# Motor impairment and adaptation in a novel non-human primate model of internal capsule infarct

**DOI:** 10.1101/2025.05.27.656471

**Authors:** Siddharth S. Sivakumar, Shashank A. Anand, Luke Brawer, Rebecca L. Mellor, Andreas Burkhalter, Harold Burton, Peter Brunner, David T. Bundy, Eric C. Leuthardt, Daniel W. Moran

## Abstract

Loss of distal hand and finger control is among the most disabling consequences of stroke. Functional outcomes are typically worse when infarcts involve subcortical white matter tracts, particularly the internal capsule, yet most preclinical stroke models target cortical regions. To address this gap, we developed a non-human primate model of internal capsule infarct using stereotactically guided endothelin-1 injections to disrupt descending fibers from the primary motor cortex hand area. Serial structural and diffusion MRI, along with histology, confirmed subcortical infarcts centered on the targeted white matter region with no apparent cortical involvement. Motor function was assessed pre- and post-infarct using a joystick-based center-out task (proximal forelimb control) and a Klüver board task (distal forelimb control). Animals exhibited variable impairments in proximal function and consistent post-infarct deficits in distal function, including reduced contralesional hand use, longer retrieval time, and increased in-well digit flexions. One animal showed mild post-infarct impairment and the smallest lesion, highlighting that this model reflects inter-individual differences in infarct size and functional outcome as seen in human subcortical stroke. In contrast, the other two animals developed a compensatory wrist-extended posture on the Klüver board task by 4 weeks post-infarct, which stabilized the hand and enabled improved digit flexion. Incorporating this behavioral adaptation into statistical models improved prediction of motor performance. The observed adaptation may have drawn on spared corticospinal output pathways, allowing animals to re-engage pre-existing motor routines to perform the retrieval. While future studies may benefit from ethologically relevant tasks to further elucidate such adaptations, findings from this study recapitulate key features of human subcortical stroke, including persistent distal motor deficits and emergence of adaptive motor strategies. By combining precise lesioning, longitudinal imaging, and detailed behavioral analysis, this model provides a translationally oriented platform for studying white matter stroke mechanisms and evaluating interventions that promote functional recovery.

## INTRODUCTION

Stroke is a leading cause of long-term disability worldwide, and motor impairment is among its most persistent and functionally limiting consequences (Thom et al., 2006; Katan & Luft, 2018). Chronic deficits in voluntary limb control, particularly of fine distal movements, often prevent return to independence and meaningful quality of life (Nakayama et al., 1994; Roby-Brami et al., 2021). Significant functional recovery beyond six months post-stroke is highly unlikely, and the extent of long-term impairment is closely tied to damage involving descending motor pathways, most notably the corticospinal tract (CST) (Stinear et al., 2007; DeVetten et al., 2010; Carter et al., 2012; Corbetta et al., 2015). Although numerous rehabilitation strategies have been explored, therapeutic options for patients with lasting motor deficits remain limited (Raffin & Hummel, 2018). One barrier to progress is the lack of preclinical models that capture the anatomical and functional complexity of stroke-induced motor impairment in humans (Edwardson et al., 2017). Most existing models, especially in rodents, rely on cortical lesions that primarily affect gray matter (Bederson et al., 1986; Carmichael, 2005; Gladstone et al., 2002), whereas the majority of strokes that result in persistent motor disability in humans involve subcortical white matter injury, including damage to the internal capsule (Corbetta et al., 2015; Kang et al., 2003; Wessels et al., 2006).

Non-human primates (NHPs) provide a critical bridge between human studies and preclinical models in lower species, offering close homology in motor system anatomy, dexterous limb function, and cortical organization (Higo, 2021). As such, NHP models are uniquely suited to address translational questions surrounding motor recovery after stroke in a mechanistic and rigorous fashion (Fisher et al., 2009). However, most prior NHP stroke models have relied on cortical infarcts or vascular occlusion techniques, which primarily affect gray matter and result in variable lesion size and location (Nudo et al., 2003; Darling et al., 2009; Zhao et al., 2014; Herbert et al., 2015). These approaches rarely produce consistent deficits in skilled forelimb use and do not replicate the focal subcortical injuries that are most often responsible for lasting motor impairment in humans (Corbetta et al., 2015; Kang et al., 2003; Higo, 2021; Kosugi et al., 2023). A reproducible model targeting descending white matter tracts, such as the posterior limb of the internal capsule, is critically needed to enable controlled investigation of motor pathway disruption and its behavioral consequences.

A prior study (Murata & Higo, 2016) demonstrated that focal infarcts could be induced in the posterior limb of the internal capsule in macaques using endothelin-1 (ET-1), a potent vasoconstrictive peptide that induces transient reduction in local cerebral blood flow through microvascular constriction. This approach established that ET-1 injections could reliably target subcortical white matter and produce sustained motor deficits, providing an important foundation for white matter stroke research in NHPs. While that study used anatomical landmarks to guide injection targeting, the present work refines this approach by aligning subject-specific structural MRI scans to both a stereotactic histological atlas and a population-averaged MRI template derived from adult rhesus macaques. This dual-atlas strategy enables more precise and reproducible targeting of internal capsule fibers, improving infarct localization while supporting structured evaluation of post-infarct behavior.

Using this refined targeting protocol, here we induced focal ET-1 infarcts in the posterior internal capsule of adult rhesus macaques, and monitored motor function and behavior over time using two task paradigms employed in several prior NHP studies. These tasks involve different aspects of forelimb motor function, with one engaging more proximal movements and the other more distal, enabling characterization of the evolving profile of impairment and recovery with greater specificity. By coupling anatomically guided infarct induction with structured behavioral assessment, this model enables reproducible targeting of subcortical motor pathways and within-subject quantification of longitudinal motor function. Additionally, this design facilitates detailed examination of how behavioral strategies evolve following injury and enables focused analysis of functional adaptation after subcortical stroke.

## METHODS

### Animals

Three adult male rhesus macaques (Mk-A, Mk-D, Mk-F; *Macaca mulatta*, 9–12 kg, 5–10 years old) were used to model a focal ischemic infarct localized to the distal forelimb region of the internal capsule. All procedures adhered to institutional and federal guidelines for the care and use of laboratory animals and were approved by the Institutional Animal Care and Use Committee of Washington University in St. Louis. Animals were housed in adjoining individual primate cages that allowed for social interaction, within a climate-controlled vivarium maintained on a 12-h light–12-h dark cycle. All experiments and procedures were conducted during the light phase. Animals and their housing environment were monitored daily by research personnel and veterinary staff to ensure health and welfare. Animals’ diet consisted of a commercial primate dry food provided twice daily, supplemented with fruits and vegetables. Fresh water was available from a drinking bottle, refreshed daily, with weekly minimum volumes (1066–1565 mL) based on individual baseline body weight. Environmental enrichment included cage-mounted commercial toys, 30 min in specialized play cages on a rotating schedule, and exposure to radio and television.

### Surgical procedures

All surgical procedures were performed under aseptic conditions and with the supervision of trained veterinary professionals in Washington University’s Division of Comparative Medicine. In each procedure, following surgical preparation, each animal was positioned prone on a level operating table and secured within a magnet-free stereotactic frame. Vital signs, monitored continuously, included heart rate, systolic and diastolic blood pressure, saturated oxygen, respiration rate, and expired carbon dioxide recorded with a gas monitoring system. A homothermic blanket system monitored and maintained core body temperature (36.6–38.3°C). No procedure required anesthesia for longer than 8 h.

#### Chamber surgery procedure

Animals were sedated with an initial injection of ketamine (6–15 mg/kg, intramuscular), and dexmedetomidine (0.01–0.03 mg/kg, intramuscular), immediately followed by atropine (0.05 mg/kg, intramuscular) and either sustained-release buprenorphine (2 mg/kg, subcutaneous) or extended-release buprenorphine (0.2 mg/kg, subcutaneous). Flunixin meglumine (0.5 mg/kg, intramuscular) and cefazolin (25 mg/kg, intramuscular) were also administered to mitigate adverse postoperative sequelae. Animals were intubated and general anesthesia was induced using inhalational administration of isoflurane gas (0.5–1.5%). Anesthesia was maintained with a mixture of 1–2% isoflurane and 50% nitrous oxide. A 22-gauge angiocatheter was inserted into the saphenous vein for continuous infusion of 0.9% sodium chloride solution (10 mL/kg/h). The scalp was shaved and washed with povidone-iodine solution in preparation for surgery.

Under aseptic conditions, a ∼15-cm midline incision on the scalp was performed, and connective tissue and temporal muscles were retracted laterally to expose the skull. Stereotactic coordinates of craniectomy centroids were determined using both a standard stereotactic atlas for Rhesus macaques (Saleem & Logothetis, 2012) and T1-weighted structural magnetic resonance imaging (MRI) scans specific to each animal, aligned to the INIA19 macaque atlas using a rigid-body transformation (FSL’s *flirt*, 6 degrees of freedom) (Rohlfing et al., 2012). On the stereotactic atlas, coordinates were identified for chamber placements both targeting the primary motor cortex and enabling vertical injection trajectories into the posterior limb of the internal capsule. These were then adjusted and validated based on each animal’s atlas-aligned scan using ear-bar zero (EBZ, defined as the midpoint along the interaural line intersected by the midsagittal plane) as a shared reference between the stereotactic and MRI atlases. Final craniectomy coordinates are reported in Table 1.

**Table 1.** Stereotactic coordinates for bilateral craniectomy centroids.

| Animal | Anterior from EBZ point (mm) | Lateral from EBZ point (mm) |
| --- | --- | --- |
| Mk-A | 18.0 | 13.5 |
| Mk-D | 17.0 | 13.5 |
| Mk-F | 17.0 | 13.5 |

The left and right parietal bones were each scored with a trephine in preparation for bilateral circular craniectomies (∼11 cm^2^ each) positioned over the central sulcus and primary motor and somatosensory cortical areas. Fifteen to seventeen holes (2.8 mm diameter) were manually drilled into the exposed skull and filled with MRI-compatible ceramic screws, forming an anchor for a skull-affixed headcap. Bilateral craniectomies were completed by removing the circular bone segments, exposing but not damaging the underlying dura mater. Sterile cylindrical acrylic chambers (16 mm inner diameter, 19 mm outer diameter) were inserted into each craniectomy site and fixed in place with dental acrylic (methyl methacrylate) applied around the chamber edges. Each installed chamber was covered with a tightly secured stainless-steel chamber cap. Additional dental acrylic, applied over the exposed skull surface, completely covered all ceramic screws to form a stable headcap. A plastic headpost was installed posteriorly on the headcap, superficial to a row of ceramic screws. At the end of the surgical procedure, atipamezole hydrochloride (0.1 mg/kg, intramuscular) was injected to reverse the effects of dexmedetomidine. Animals were then transferred to their home cages and monitored during emergence from anesthesia and recovery. Veterinary staff provided supplemental postoperative care as needed.

#### Infarct procedure

Lyophilized human-derived ET-1 (Sigma, E7764) was dissolved in diluted acetic acid to a final concentration of 0.5 µg/µL. The solution was filtered using a Spin-X centrifugal filter, aliquoted into 0.5-mL microcentrifuge tubes in 10-µL portions, and stored in ice at −20°C. Each 10-µL aliquot was designated for a single penetration, comprising a triplet of injection sites aligned along the superior–inferior axis. In each animal, ET-1 injections were localized to the posterior limb of the internal capsule corresponding to descending white matter tracts from the M1 hand area. The left hemisphere was selected because all three animals exhibited a strong right-hand preference as determined by spontaneous use of the right hand in >65% of Klüver board task trials (see “Klüver board task”). Target injection sites were identified using a macaque stereotactic atlas (Saleem & Logothetis, 2012), anatomical data from a tracer study that identified motor fiber locations in the posterior limb of the internal capsule in NHPs (Morecraft et al., 2002), and MRI atlas-aligned parcellations of animal-specific T1- and T2-weighted structural scans (see “Magnetic resonance imaging procedures”). Injection sites were also chosen such that a direct vertical trajectory to the target could be achieved through the exposed dura of the left cortical chamber.

Surgical preparation, anesthesia, and sterile technique for the infarct procedure followed the same protocol as described for the chamber surgery. To aid precise infarct targeting, a custom guide tube was fabricated by cutting the tip off a 22-gauge needle and smoothing the bevel from the cut edge using a rotary tool. To prevent slippage during use, a small metal ball was soldered at the midpoint of the guide tube under hydrochloric acid flux. The guide tube was used to position the 26-gauge needle of a 10-µL microsyringe, which was mounted to a stereotactic frame and advanced vertically (inferiorly) through the guide tube, penetrating the dura, cortex, and entering the internal capsule. At each of the three target sites per penetration, 3.0–3.3 µL of the ET-1 solution was injected at a rate of 0.5–1 µL/min (site coordinates reported in Table 2). Following each injection, the needle was held in place for 10–15 min to allow the solution to diffuse and minimize backflow along the needle track. The needle was then retracted along the superior direction to the next injection depth. After completing all three injections for a given penetration, the needle was fully withdrawn and repositioned for the next penetration site.

**Table 2.**
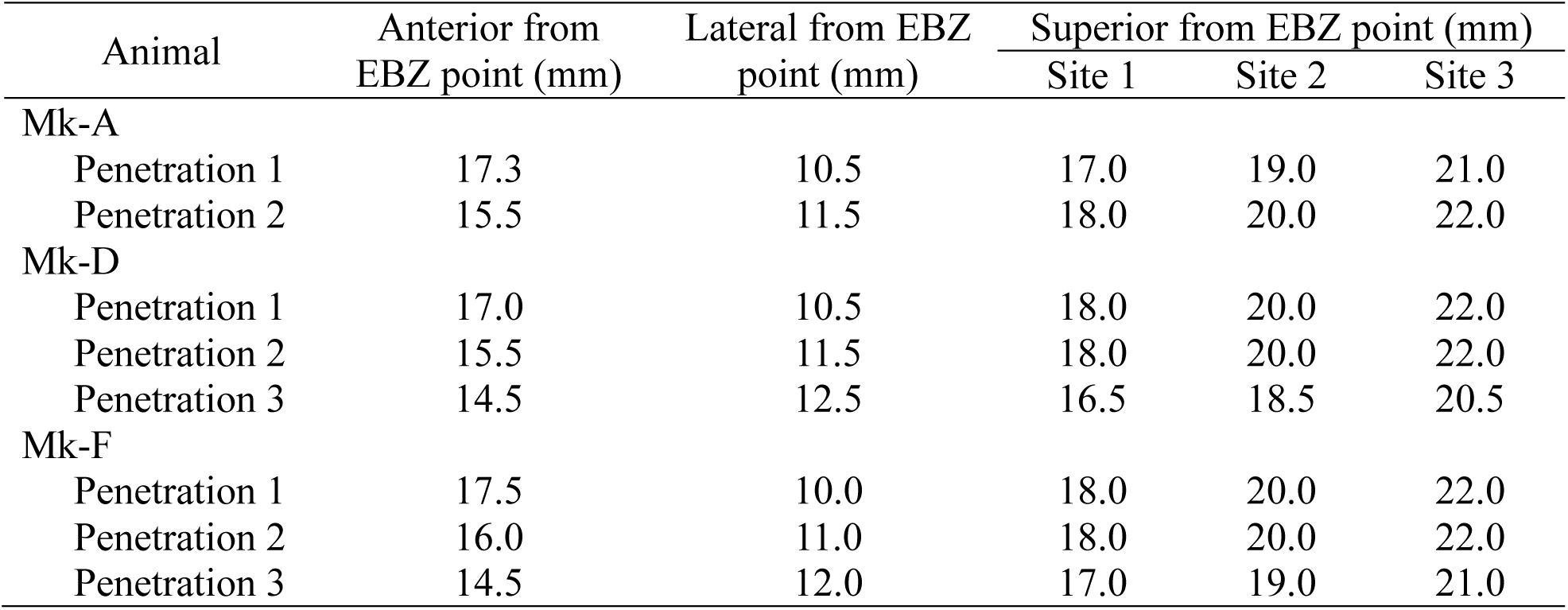
Stereotactic coordinates for left internal capsule ET-1 injection sites.

Upon completion of the final injection, the cortical chamber was inspected to ensure no bleeding or fluid leakage from the penetration sites, which were subsequently cleaned with sterile saline. A sterile stainless-steel cap sealed the chamber. Anesthesia was discontinued, the animal was returned to its home cage, and was monitored during recovery for any overt motor deficits by researchers and veterinary staff. Routine enrichment was continued for each animal post-infarct. However, the cage-mounted foraging board, which required fine and dexterous finger movements, was intentionally withheld during the post-infarct period to minimize unsupervised rehabilitation of distal motor function outside of the Klüver board task. This increased the likelihood that observed changes in task performance reflected spontaneous functional recovery rather than task-irrelevant motor learning.

#### Post-surgical validation of infarct location

Approximately 1 week following infarct induction, structural MRIs were acquired to validate infarct location. Each animal’s T1-weighted image was aligned to the INIA19 macaque stereotactic atlas (Rohlfing et al., 2012), and the ear-bar zero (EBZ) was identified as a reference point. The centroid (3D mean) of the injection trajectories was then calculated and compared against an internal capsule mask to assess the accuracy of injection targeting.

### Behavioral procedures

Two behavioral paradigms with extensive prior use in NHPs were selected to assess motor function at proximal and distal levels of the forelimb. A joystick-based center-out task assessed primarily proximal control, while a Klüver board task assessed fine distal movements. Together, these paradigms enabled task-specific evaluation of motor control and functional recovery during the pre- and post-infarct periods.

#### Center-out task with joystick control

As described in prior studies by our lab and others (Georgopoulos et al., 1986; Moran & Schwartz, 1999; Pearce & Moran, 2012; Chan & Moran, 2006), animals were trained using operant conditioning to perform joystick movements, controlling a cursor on a screen in a standard 2D center-out task (Figure 1A). All movements were performed by the animal while it was seated comfortably in a primate chair, grasping a chair-mounted, cylindrical joystick (18 mm diameter, 22 mm length) positioned at elbow height and directly in front of the animal’s torso. Joystick position was mapped directly to cursor (*x*, *y*)-position on the screen, with 1 cm of joystick displacement corresponding to 6 cm of displayed cursor movement on the screen. Two video cameras recorded the animal’s head, arms, and torso. An experimenter in a separate room continuously monitored animal behavior by video and audio to ensure the animal’s comfort and welfare. Cursor position was continuously recorded throughout each trial at 60 Hz and synchronized to 30-Hz video recordings using the BCI2000 software platform (Schalk et al., 2004).

**Figure 1.**
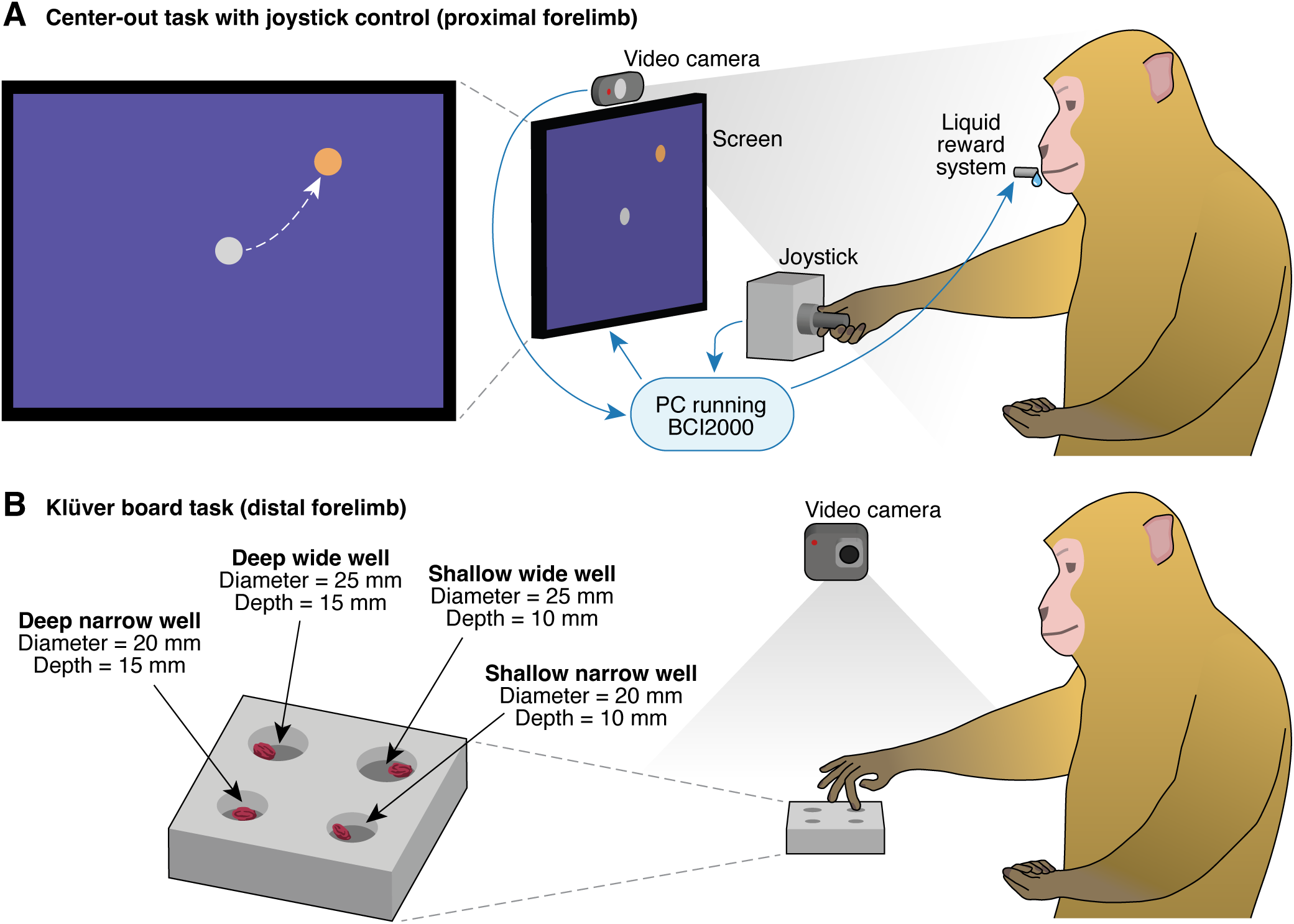
Motor function assessment tasks. **A.** Center-out task with joystick control. This task was used primarily to train and evaluate proximal forelimb control, in addition to some distal forelimb skill required to grasp the cylindrical joystick handle (diameter 18 mm, length 22 mm). Animals controlled an on-screen cursor using the joystick, sequentially moving it to a central target and then to one of eight peripheral targets selected pseudorandomly within trial blocks. Trials were successful if the animal maintained cursor position within the central and outer targets for randomized hold durations, after which a liquid reward (2–4 mL) was delivered. Cursor position data (60 Hz) and video recordings of the animal’s head and forelimb movements (30 Hz) were synchronized and acquired simultaneously using the BCI2000 software platform. **B.** Klüver board task. This task was used to train and evaluate skilled distal forelimb movements, particularly controlled digit flexion and grasping during reach-and-retrieval movements. Monkeys retrieved a single small morsel of food (either a pellet or a dry raisin) from each of four wells ranging from 20–25 mm in diameter and 10–15 mm in depth. Forelimb movements during reaching and retrieval were recorded at 120 Hz by two tripod-mounted cameras (only one camera shown), each laterally positioned and angled downward at 45° toward the center of the Klüver board. An acrylic divider (not pictured) restricted animals to use their right forelimb during approximately half of the trials; forelimb choice was unrestricted during remaining trials.

Throughout each trial, the cursor remained visible on screen. The cursor was circular, with a diameter equal to that of circular central and outer targets (2–4 cm). A flowchart illustrating the sequence of trial events and possible outcomes is provided in Figure 4A. Each trial began when the central “start” target appeared, prompting the animal to capture this target by moving the cursor so that at least part of it overlapped with the central target region. If the animal failed to capture or hold the cursor within the central target within a preset maximum duration or did not maintain position for the randomized hold-A period (318–530 ms), the trial was classified as not attempted.

If the animal successfully held the cursor within the central target for the required hold-A duration, an outer target subsequently appeared on screen. Outer targets were positioned at one of eight possible peripheral locations equally spaced at 45° intervals around the central target, with 0° defined as directly to the right. Target selection was arranged into trial blocks, each consisting of 16 trials in which each of the eight outer targets appeared exactly twice per block, selected without replacement and with uniform probability. Although the complete task sequence for each experimental session comprised 512 trial blocks, no more than 30% of this sequence was completed in a single session.

Following the appearance of the outer target, a randomized delay period (205–410 ms) occurred, during which the animal was required to maintain cursor position steadily on the central target. Premature cursor movement toward the outer target, before the disappearance of the central target, resulted in the trial being classified as attempted but not successful (Figure 2A). At the end of this delay period, the central target disappeared, serving as a “go” cue signaling the animal to rapidly move the cursor outward toward the outer target. Movement initiation (reaction time) and execution (movement time) were both constrained within predefined time windows (61–5000 ms and ≤5000 ms, respectively).

**Figure 2.**
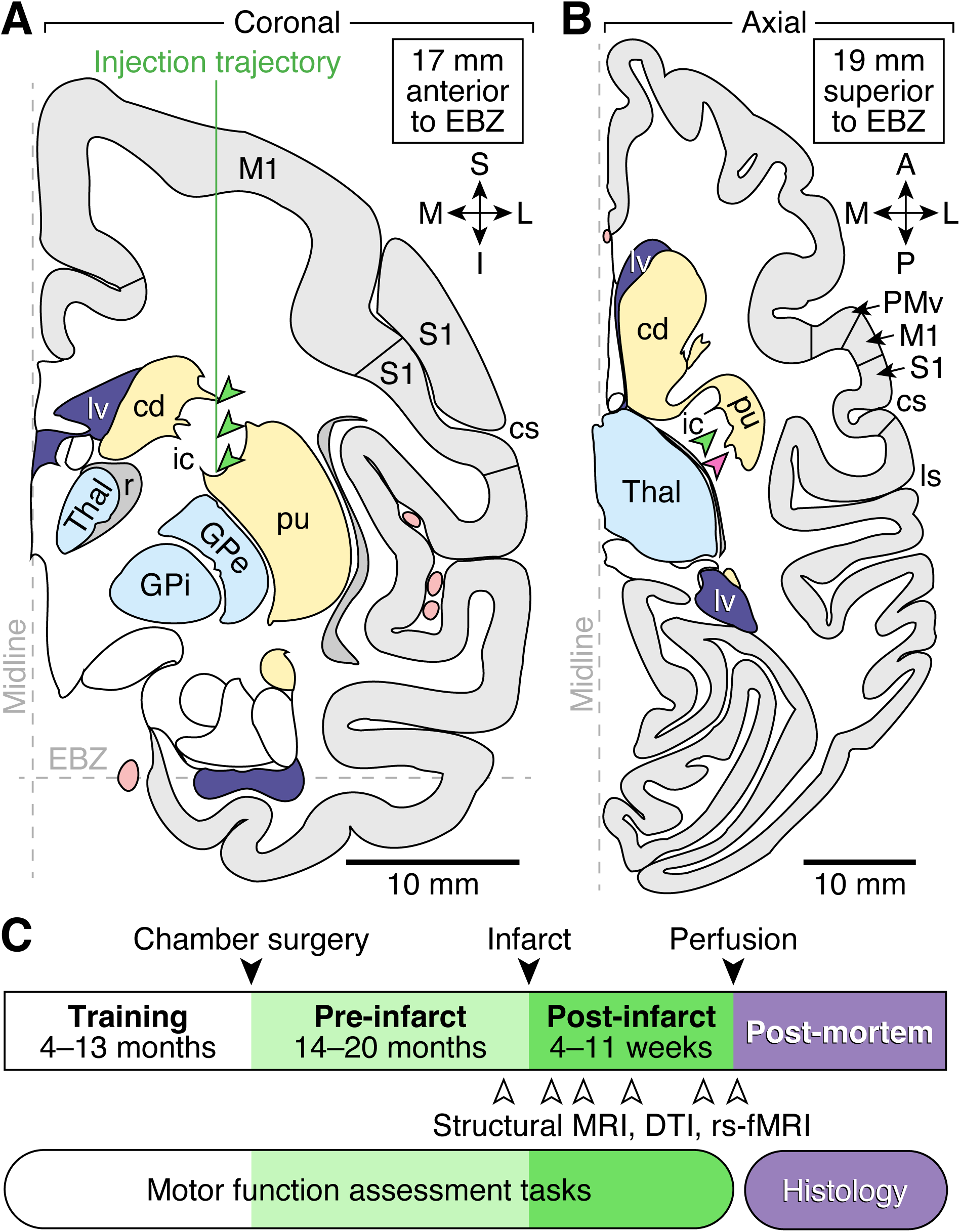
Infarct targeting and longitudinal study design. **A.** A vertical injection trajectory for endothelin-1 (ET-1) was facilitated by a 16-mm-wide cortical chamber positioned over the left primary motor cortex (M1). Stereotactic coordinates for needle penetration were determined using a standard Rhesus macaque stereotactic atlas and aligned with animal-specific T1-weighted structural MRI scans. For each penetration, ET-1 (3.0–3.3 µL of a 0.5 µg/µL solution) was injected at each of three vertically spaced sites (green arrowheads). **B.** Injection trajectories for a single animal (Mk-A). Green and magenta arrowheads indicate two separate penetrations; the green arrowheads correspond to the injection trajectory shown in **A**. Diagrams in **A–B** were adapted from Saleem and Logothetis (2012). Abbreviations: cd, caudate nucleus; cs, central sulcus; EBZ, ear-bar zero; GPe, external segment of globus pallidus; GPi, internal segment of globus pallidus; ic, internal capsule; ls, lateral sulcus; lv, lateral ventricle; M1, primary motor cortex; PMv, ventral premotor cortex; pu, putamen; r, reticular nucleus; S1, primary somatosensory cortex; Thal, thalamus. Anatomical orientation markers: M, medial; L, lateral; S, superior; I, inferior; A, anterior; P, posterior. **C.** Animals were trained for several months prior to surgery to perform the two motor assessment tasks described in Figure 1, continuing these tasks post-infarct 1–5 days per week throughout the entire study. Craniectomies for cortical chamber installation marked the onset of the pre-infarct period. Structural, diffusion, and functional MRI scans were acquired 1–2 months before infarct induction, and again at 3–4 time points following infarct. On the day of euthanasia and perfusion, additional MRI sequences were obtained from the animal’s head. Extracted brain specimens underwent histological sectioning and staining.

Once the central target disappeared, three trial outcomes were possible. If the cursor remained stationary at the center of the screen (within the original boundary of the central target despite its disappearance), the trial was categorized as not attempted, as this typically indicated that the animal was no longer actively holding the joystick, which automatically returned to a neutral center position when released. If the cursor moved outward but either failed to capture the outer target or failed to hold it for the randomized hold-B duration (295–530 ms), the trial was categorized as attempted but not successful. Finally, trials in which the animal both captured and maintained cursor position within the outer target region for the required hold-B period were categorized as attempted and successful. Note that on each trial, the hold-A, delay, and hold-B periods were independently randomized from uniform distributions (ranges specified above) to minimize anticipatory timing strategies in favor of cue-driven movement initiation.

#### Klüver board task

This task was adapted from previous investigations of fine motor control of the distal forelimb in NHPs (Won et al., 2020; Moore et al., 2019; Nudo et al., 1996; Friel & Nudo, 1998; Plautz et al., 2000; Pizzimenti et al., 2007). A transparent polycarbonate block (10 × 10 × 2.5 cm) containing four wells of varying dimensions was affixed to a horizontal stand that could be repositioned by the experimenter (Figure 1B). The four wells varied systematically in diameter and depth: shallow wide well (SW; 25 mm diameter, 10 mm depth), shallow narrow (SN; 20 mm diameter, 10 mm depth), deep wide (DW; 25 mm diameter, 15 mm depth), and deep narrow (DN; 20 mm diameter, 15 mm depth). On each trial, one small morsel of food (pellet or dry raisin) was placed in each well, and the board was positioned within the animal’s reach, aligned to the same plane as the animal’s waist and ∼20–30 cm in front of its torso.

On approximately half of the trials, a vertically oriented, T-shaped acrylic divider was positioned to permit only right forelimb use; the divider featured a segment extending toward the animal with a 10-cm-diameter hole on the right side. During the remaining trials, the divider was removed, allowing unrestricted forelimb choice. Two tripod-mounted video cameras (GoPro Hero8 Black) positioned laterally (∼1 m away), each angled downward at 45° toward the center of the Klüver board, recorded reaching and retrieval movements at 120 Hz. Each trial block consisted of one reach to each of the four wells. A trial was considered successful if the animal retrieved and transferred the food item to its mouth. On days when the Klüver board task was performed, each animal attempted 4–20 trial blocks. All animals learned to perform the task successfully within 5 days of training.

### Behavioral data analyses

Quantitative analyses of both behavioral paradigms were performed at the trial level using either generalized linear mixed models (GLMMs) or linear mixed models (LMMs). These models included fixed effects for infarct status and other task-relevant predictors, along with random effects to account for per-session variability. This approach accounted for trial-wise variability and within-subject changes over time, enabling more rigorous characterization of the effects of infarct and other factors on motor performance. All behavioral analyses were conducted using MATLAB (version R2024b; MathWorks, Natick, MA) or R (version 4.4.2; R Core Team, Vienna, Austria).

#### Center-out task analyses

The origin (*x* = 0, *y* = 0) was defined as the center of the screen, coinciding with the center of the central target. Cursor *x*- and *y*-position time series were each normalized to the range [−0.5,0.5] for all analyses. Trial epochs were defined from the time of central target appearance until either a successful hold at the outer target or occurrence of any failure condition (see “Center-out task with joystick control”). For the purpose of determining if the animal attempted movement after the central target disappeared, a trial was classified as attempted if the cursor’s radial position *r* = √*x*^2^ + *y*^2^ exceeded the 5th percentile threshold of session-wide *r* values. For each successful trial, the movement epoch was defined to begin at the first time *r* exceeded 5% of the distance between the origin and the center of the outer target, and to end when the cursor first overlapped the outer target (i.e. when the distance between cursor center and outer target center fell below one cursor diameter). This epoch definition was chosen to minimize the effects of pre-movement planning and reaction delays in the analysis of volitional movement. For successful trials, movement duration was calculated as the time of this epoch (in seconds), and path length as the sum of Euclidean distances between successive (*x*, *y*) samples over the movement epoch. Prior to statistical modeling, path-length values were converted to length on screen by applying a 24-cm scaling factor.

To assess the effect of infarct status on two trial-level binary outcomes—whether a trial was attempted, and whether an attempted trial was successful—GLMMs were fit separately for each animal. Each model used a logistic link function to model the probability of the binary outcome as a function of infarct status, with session included as a random intercept to account for baseline variability across sessions. Each model was defined as

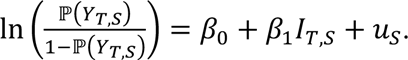

Here, 𝑌_*T*,*S*_ ∈ {0,1} indicates either trial attempt (1) or trial success conditional on attempt (1) for trial *T* in session *S*; 𝐼_*T*,*S*_ ∈ {0,1} indicates infarct status (1 for post-infarct); the fixed effects *β*_0_ and *β*_1_ represent the log-odds of the outcome in the pre-infarct condition and the change in log-odds associated with the post-infarct condition, respectively; and 𝑢_*S*_ ∼ 𝒩(0, 𝜎^2^) is a session-specific random intercept accounting for baseline differences in outcome probability across sessions. Importantly, the model for 𝑌_*T*,*S*_ = attempts included all trials, whereas the model for 𝑌_*T*,*S*_ = successes included only attempted trials.

To evaluate the effect of infarct status on path length and movement duration on successful trials, separate LMMs were fit for each animal using log-transformed trial-wise values as the outcome variable. These models differed from the GLMMs described above in that they assumed normally distributed residuals and modeled the logarithm of the continuous outcome variable rather than a probability. The log-transformation was applied to account for right-skewed distributions observed in both metrics. Infarct status was included as a fixed effect, and session was modeled as a random intercept. Each model took the form

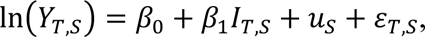

where 𝑌_*T*,*S*_ represents either path length or movement duration for trial *T* in session *S*; 𝐼_*T*,*S*_ ∈ {0,1} indicates infarct status; *β*_0_ and *β*_1_ denote the fixed effects for the pre-infarct baseline and post-infarct shift; 𝑢_*S*_ ∼ 𝒩(0, 𝜎^2^) is a session-specific random intercept; and 𝜀_*T*,*S*_ ∼ 𝒩(0, 𝜏^2^) is the residual error term. Both LMMs included only trials that were successful.

The numbers of trials and sessions included for all models described in this section are summarized in Table 3. In Figure 4B–C (Mk-A), Figure S1A–B (Mk-D), and Figure S2A–B (Mk-F), the left side of each panel shows either the proportion of trials attempted per session or the proportion of successful trials among attempted trials (dots), along with the mean rate across sessions (bars). In Figure 5B–C (Mk-A), Figure S1C–D (Mk-D), and Figure S2C–D (Mk-F), the left side of each panel shows the distributions of trial-level values, pooled across all sessions included. Estimated marginal means were derived from each model along with 95% confidence intervals, as shown on the corresponding right side of each figure panel referenced above. These values were computed from each model’s fixed-effect estimates using the *predict* function in MATLAB, by generating predicted probabilities for the pre-infarct and post-infarct conditions, designating a representative session as a placeholder for the random effect. This approach reflects the uncertainty in the estimated probabilities while holding session-level variation constant.

**Table 3.**
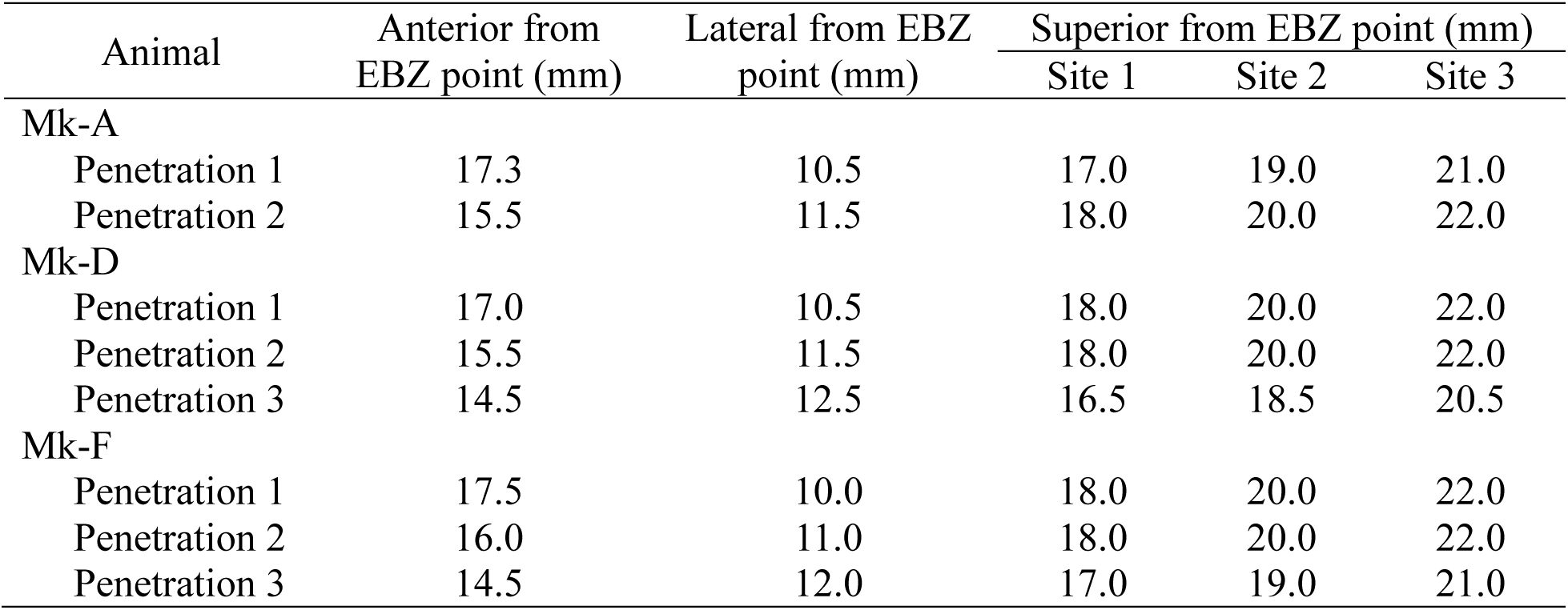
Summary of sample sizes included in center-out task models.

For center-out task kinematic analysis, the raw cursor trajectory for a successful trial was defined as the (*x*, *y*)-time series beginning at the sample when the central target disappeared and ending at the final sample of the hold-B period. This epoch definition was chosen to adequately capture both movement initiation at the central target and terminal stabilization at the outer target. Raw cursor trajectories are colored by outer target in Figure 5A.

To estimate where future cursor trajectories would be likely to fall, 95% prediction regions were estimated from variability along the local normal direction of the mean trajectory, across successful trials in a session. For each outer target, raw trial-wise (*x*, *y*) cursor trajectories were first linearly interpolated to 1000 samples. At each sample 𝑡 (excluding the final sample), the mean trajectory 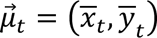 was computed from the interpolated trajectories (and is shown as a thick black trace in Figure 5A,D), and the unit tangent vector was computed as 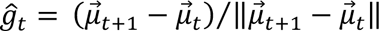. The corresponding unit normal vector 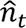 was defined as 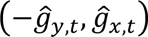. The deviation vector of the interpolated trajectory on trial *T* from the mean trajectory was defined as 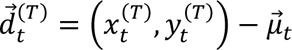. Each deviation vector 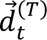 was projected onto 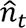, and the standard deviation 𝜎_𝑡_ of these scalar projections was computed across trials for each sample 𝑡. 95% prediction region endpoints along the normal direction 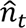 were then calculated as 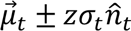, where *z* = ϕ^−1^(0.975) ≈ 1.96. The standard deviation—not the standard error—was used because the goal was to estimate variability across individual future trajectories, not uncertainty of the mean. The resulting left and right endpoint sequences were each smoothed across samples using a moving average filter with a window size of 200 samples, to stabilize the region boundaries.

To avoid including reversals in radial displacement that occurred after the outer target was reached, each mean trajectory was projected onto the fixed radial direction associated with its target (i.e., the vector from the origin to the center of the outer target). The final 100 samples of this 1D projection were scanned for the earliest occurrence of three or more consecutive negative changes, which would indicate a consistent reversal in radial movement. The mean trajectory and its associated prediction region were then truncated at the last sample before this reversal or retained in full (𝑡_234_ = 1000) if no such reversal was detected. To avoid sharp boundary edges near the outer target, a linear ramp-down was applied over the final 20 samples of the left and right endpoint sequences. Each endpoint was linearly scaled toward the corresponding mean trajectory position, such that the region boundary smoothly collapsed to the mean by the final sample. The resulting 95% prediction region (Figure 5D) estimates the spatial area within which 95% of future cursor trajectories are expected to fall.

#### Klüver board task analyses

Video recordings from each session were manually scored by a trained experimenter who was blinded to both the date of the session and the identity of the animal. For each trial, the following variables were recorded: the trial index within the session, the hand primarily used for retrieval, whether hand use was restricted to the right forelimb via an acrylic divider, and the identity of the target well, classified as SW, SN, DW, or DN. Trial success was noted based on whether the animal successfully transferred the food to its mouth. Two behavioral metrics were predefined to quantify task difficulty: retrieval time and the number of in-well digit flexions. Retrieval time was defined as the duration from the onset of volitional forelimb movement toward the target well to the moment the food item contacted the animal’s mouth. This duration was calculated in video frames and converted to seconds. In-well digit flexions were defined as discrete flexion movements at the distal interphalangeal (DIP) joint of a single digit while it was positioned inside the target well. The goal of each flexion was to press the food item between the fingertip and the interior wall of the well, allowing the item to be rolled upward along the wall to facilitate grasp. These movements were almost exclusively executed by digit II, with <2% of trials involving digits I or III. Sample sizes for all Klüver board analyses, including the number of sessions and trials by animal and hand used, are reported in Table 4.

**Table 4.**
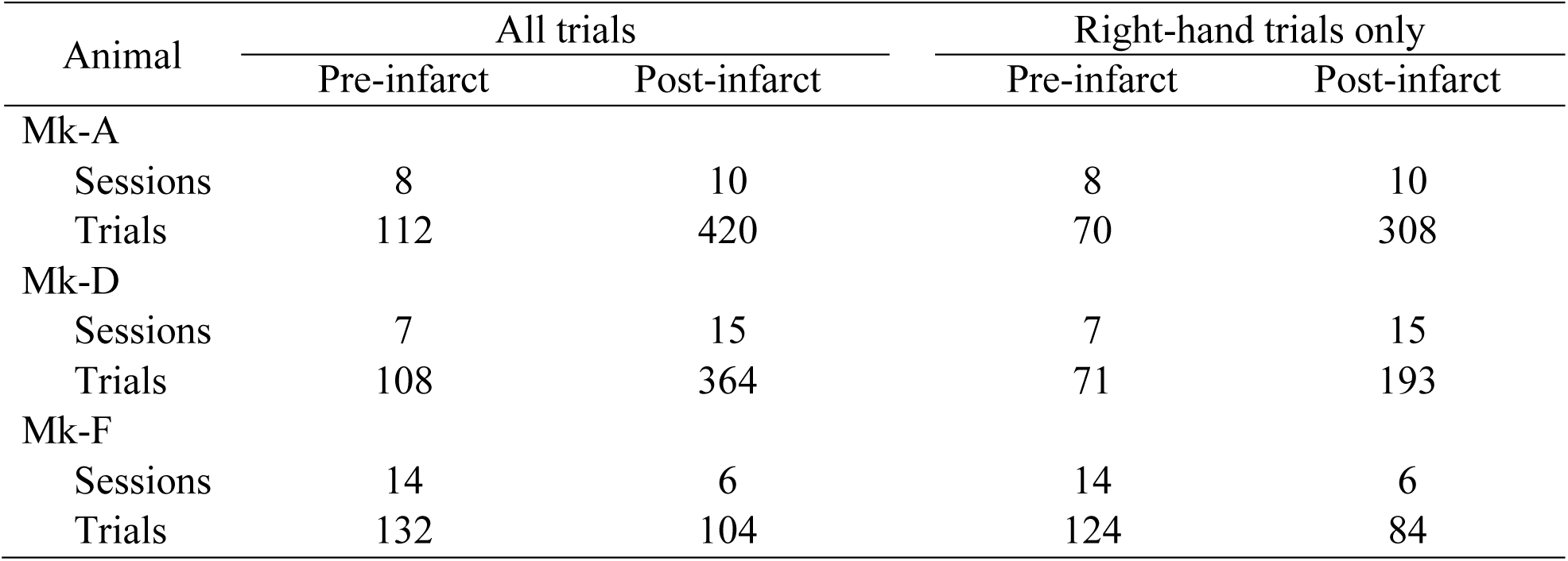
Summary of sample sizes for Klüver board task data.

Following initial video scoring and further behavioral observation, it became evident that two animals (Mk-A and Mk-D) altered their strategy for performing the Klüver board task in the post-infarct period, as described in the Results section. To quantify this behavioral adaptation, we identified two trial-level metrics that captured key changes in hand posture: wrist extension and digit-II extension. These metrics were evaluated using individual trial clips segmented from raw video recordings to isolate hand movements near the Klüver board. Each clip was then scored by a trained experimenter who was blinded to the date of the trial, the trial index within the session, and the identity of the animal. Wrist extension was defined as present on a given trial if, at the time digit II contacted the target well, the wrist was extended in a posture that resulted in the flexed digits III–V pressing firmly against the surface of the Klüver board (Figure 6F, right). Digit II was considered extended if, at any time while the hand was above or making contact with the Klüver board, the DIP joint of digit II was fully or nearly fully extended (>160°). These binary adaptation metrics were chosen for their simplicity and for their ability to capture task-relevant biomechanical changes post-infarct. Specifically, they reflected both impaired digit extension and postural adjustments that typically improved in-well DIP flexion by allowing digit II to enter the well at a more board-orthogonal angle.

To assess the effects of infarct on task difficulty, and to additionally evaluate the effects of behavioral adaptation, GLMMs were fit to retrieval time and in-well digit flexions from right-hand trials pooled across all animals. For retrieval time, a Gamma distribution with a log link function was used to account for the right-skewed, continuous outcome. For in-well digit flexions, a Poisson distribution with a log link was used, as this metric represents discrete count data. First, models were fit without adaptation metrics, including fixed effects for infarct status and well type, and random intercepts and slopes for days since infarct within animal:

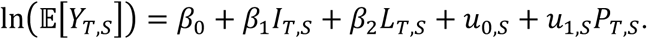

Next, models with adaptation metrics added fixed effects for wrist extension and digit-II extension:

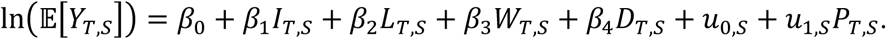

For the above model definitions, 𝑌_*T*,*S*_ denotes the observed outcome on trial *T* in session *S*; 𝐼_*T*,*S*_ ∈ {0,1} indicates infarct status; 𝐿_*T*,*S*_ denotes well type, treated as a four-level categorical variable (SW, SN, DW, DN) and encoded using indicator variables with DN as the reference level; 𝑊_*T*,*S*_ ∈ {0,1} indicates the presence of wrist extension; 𝐷_*T*,*S*_ ∈ {0,1} indicates the presence of digit-II extension; and 𝑃_*T*,*S*_ indicates days post-infarct. The fixed-effect coefficients *β*_0_ through *β*_4_ represent the intercept and the effects of infarct status, well type, wrist extension, and digit-II extension, respectively. The terms 𝑢_0,*S*_ and 𝑢_1,*S*_ represent animal-specific random intercepts and slopes with respect to 𝑃_*T*,*S*_. A fixed effect for hand restriction (restricted vs. unrestricted) was initially used but excluded from final models due to lack of significant improvement in model fit. In Figure 6D–E, the change in the difficulty metric between pre- and post-infarct periods corresponds to the estimated coefficient *β*_1_, and the associated *p*-value reflects the significance of that coefficient based on a Wald 𝑡-test.

To test the added explanatory value of behavioral adaptation, a likelihood-ratio test was performed to compare each model without adaptation metrics (nested model) to the corresponding model with adaptation metrics (full model). Model fit was further quantified using an estimated 𝑅^2^ value (Figure 6H), representing the proportion of variance in the observed outcomes explained by the model. This was calculated as the squared Pearson correlation between model-predicted and observed responses.

### Magnetic resonance imaging procedures

Animals were initially anesthetized with ketamine (4 mg/kg, intramuscular) and dexmedetomidine (0.01 mg/kg, intramuscular), immediately followed by atropine (0.05 mg/kg, intramuscular). Subsequently, stainless-steel chamber caps were replaced with MRI-compatible plastic caps. Animals were then intubated, and anesthesia was maintained using 0.5–1.5% isoflurane delivered through a custom MRI-compatible system. Animals were positioned head-first and supine in the scanner. Imaging was performed using a 3T Siemens PRISMA system (Siemens, Erlangen, Germany) equipped with a 24-channel receive-only coil array (Takashima Seisakusho Co., Rogue Research, Montreal, Canada) housed in a split coil design.

T1-weighted structural images were acquired using a 3D magnetization-prepared rapid acquisition gradient echo (MPRAGE) sequence (Mugler III & Brookeman, 1991) with the following parameters: 0.5 mm isotropic voxel resolution, field of view (FOV) 128 × 128 × 64 mm, matrix 256 × 256, 128 slices per slab, transversal orientation, anterior-to-posterior readout direction, 25% slice oversampling, 15% phase oversampling, two averages, repetition time (TR) = 3200 ms, echo time (TE) = 562 ms, inversion time (TI) = 800 ms, bandwidth = 140 Hz/pixel, no fat suppression, no GRAPPA acceleration, turbo factor 160, and no prescan normalization.

T2-weighted structural images were acquired using a Sampling Perfection with Application optimized Contrast using different angle Evolutions (SPACE) sequence (Mugler et al., 2000) with the following parameters: 0.5 mm isotropic voxels, FOV 128 × 128 × 64 mm, matrix 256 × 256, 128 slices per slab, transversal orientation, anterior-to-posterior phase-encoding direction, 25% slice oversampling, 15% phase oversampling, two averages, TR = 3200 ms, TE = 562 ms, echo train duration = 1453 ms, bandwidth = 383 Hz/pixel, no fat suppression, no GRAPPA acceleration, turbo factor 314, and no prescan normalization.

Diffusion-weighted images were acquired using a 2D spin-echo echo-planar imaging (EPI) sequence with a monopolar diffusion scheme and 𝑏-values of 1000 and 2000 s/mm^2^. Two scans were acquired with reversed phase-encoding directions (anterior-to-posterior and posterior-to-anterior). Imaging parameters included: 1.5 mm isotropic voxels, FOV 138 × 138 × 60 mm, matrix 92 × 92, 40 interleaved slices, GRAPPA acceleration factor = 2, TR = 5000 ms, echo spacing = 1.06 ms, bandwidth = 1046 Hz/pixel, no multiband acceleration, partial Fourier phase encoding off, and no prescan normalization. Diffusion encoding included 66 noncollinear directions over a half sphere and 12 𝑏 = 0 volumes (six acquired before and six interleaved during each scan).

Spin-echo EPI images for 𝐵_0_ field map estimation were acquired using reversed phase-encoding directions (anterior-to-posterior and posterior-to-anterior) with the following parameters: FOV 138 × 138 × 60 mm, 1.5 mm isotropic voxels, transversal orientation, interleaved slices, TE = 88.60 ms, echo spacing = 1.06 ms, bandwidth = 1046 Hz/pixel, no partial Fourier, no fat suppression, and no prescan normalization.

At the conclusion of imaging, stainless-steel chamber caps were reinstalled, and atipamezole hydrochloride (0.1 mg/kg, intramuscular) was administered to reverse the effects of dexmedetomidine.

To estimate infarct volume at 1 week post-infarct, hyperintense voxels were identified by applying a fixed threshold to T2-weighted images in raw scanner units. Thresholds were selected individually for each animal based on visual inspection of infarct-associated signal at the ET-1 injection site. Voxel counts above the selected threshold were multiplied by voxel volume (0.125 mm^3^) to estimate infarct size. Threshold values and corresponding infarct volume estimates are reported in Table 5.

**Table 5.** Animal-specific infarct volume estimates at 1 week post-infarct from T2-weighted MRI.

| Animal | Estimated infarct volume (mm<sup>3</sup>) | T2-weighted MRI threshold used (raw scanner units) |
| --- | --- | --- |
| Mk-A | 164.25 | ≥120 |
| Mk-D | 914.5 | ≥89 |
| Mk-F | 139.75 | ≥80 |

If large differences in baseline voxel intensity were observed across scans, mean intensity was equalized by adding or subtracting a fixed integer to all voxels within that scan. This normalization was performed independently for each modality (T1, T2, and DTI) and applied only for visualization purposes in Figures 3A, 7A–B, S3, and S4.

**Figure 3.**
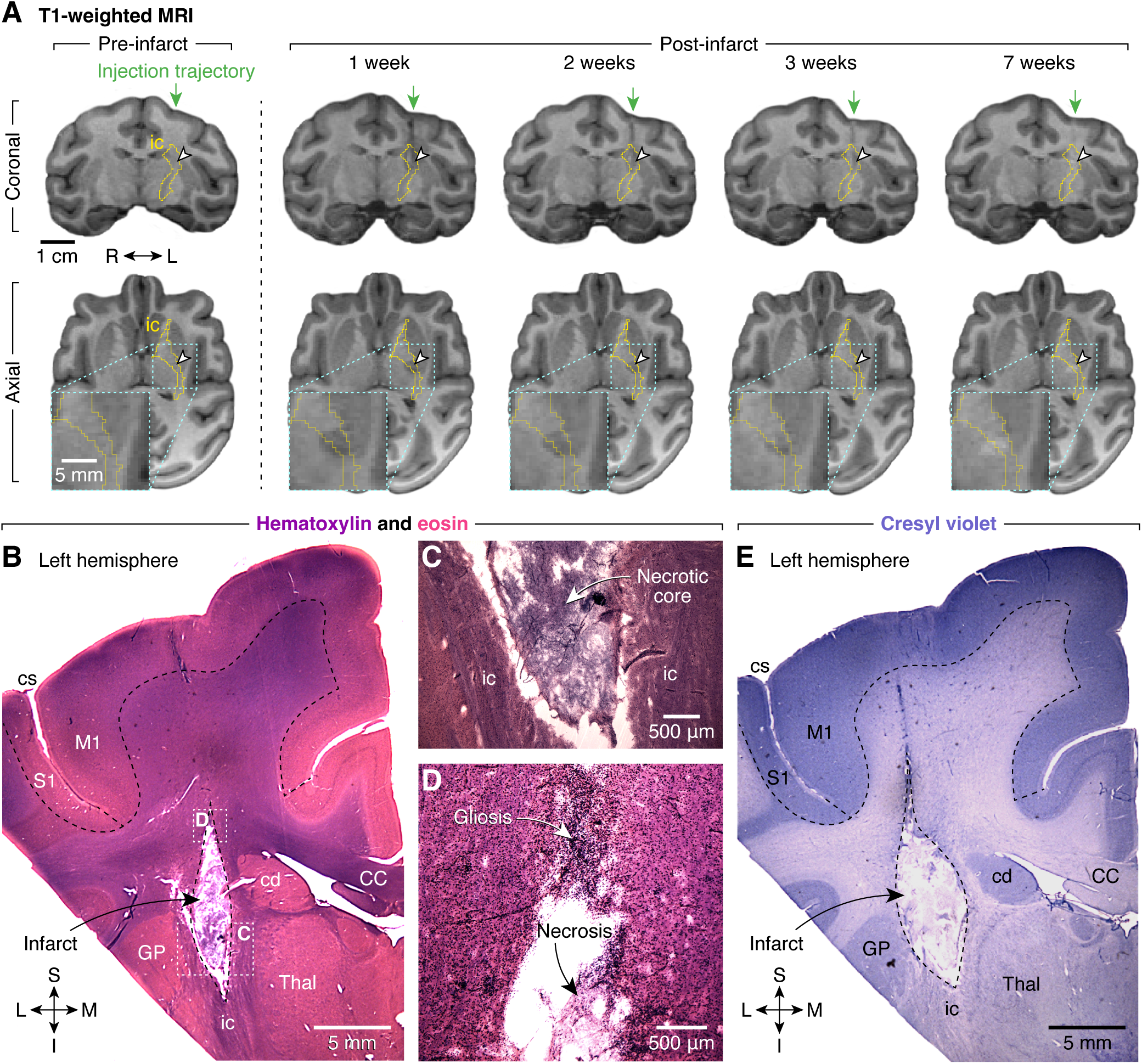
T1-weighted MRI and histological characterization of ET-1-induced internal capsule infarct in an exemplar animal (Mk-A). **A.** Longitudinal T1-weighted MRI scans illustrating infarct evolution in the internal capsule. Each column shows coronal (top row) and axial (bottom row) slices at consistent atlas-aligned coordinates. Internal capsule (ic) is outlined in yellow; ET-1 injection centroid is marked by a white arrowhead (planned in pre-infarct, actual in post-infarct images). Coronal slices include a green arrow denoting the vertical injection trajectory. Axial slices include a bottom-left inset enlarging the region near the injection site. Anatomical orientation marker indicates right (R) and left (L) hemispheres. Pre-infarct scan shows normal T1 signal near the injection site. Post-infarct scans were acquired at 1, 2, 3, and 7 weeks. Hypointense voxels near the injection centroid are evident at 1–3 weeks and diminish over time; by 7 weeks, the region appears hyperintense, consistent with chronic infarct. **B.** Hematoxylin-and-eosin-stained coronal section through the left hemisphere, showing a focal infarct confined to white matter fibers of the internal capsule. Note the preservation of adjacent subcortical structures such as the caudate nucleus, globus pallidus, and thalamus. Regions marked by white dotted rectangles are enlarged in **C–D**. **C.** High-magnification image of the infarct core, characterized by tissue pallor, vacuolization, loss of normal white matter architecture, reduced cellularity, and fragmented eosinophilic debris, consistent with infarct-related necrosis. **D.** Peri-infarct region demonstrating increased cellular density, indicative of reactive gliosis. **E.** Cresyl violet staining further confirms cellular loss and pallor within the infarct core of the internal capsule, consistent with white matter degeneration. Abbreviations in **B**, **C**, and **E**: CC, corpus callosum; cd, caudate nucleus; cs, central sulcus; GP, globus pallidus; ic, internal capsule; M1, primary motor cortex; S1, primary somatosensory cortex; Thal, thalamus. Anatomical orientation markers in **A** and **D**: M, medial; L, lateral; S, superior; I, inferior. Images in all panels are from a single animal (Mk-A).

### Histological procedures

#### Perfusion and sectioning

Anesthetized animals were catheterized in the saphenous vein using a 22-gauge angiocatheter. Heparin (10,000 units in 1 mL, intravenous) was administered to achieve systemic anticoagulation over 5 min. Sodium pentobarbital (150 mg/kg, intravenous) was then injected and cessation of cardiac and respiratory activity was subsequently confirmed by auscultation. Animals were placed supine on a necropsy table, and a midline incision (∼30 cm) was made from the clavicle to below the ribcage. The lateral aspects of the ribs were transected bilaterally, and the sternum was separated from the clavicles. The sternum and anterior ribs were then removed en bloc to expose the mediastinum. After isolating the heart and removing the pericardium, the right atrium was incised as a drainage site, and the left ventricle was cannulated. The circulatory system was flushed with 2 L of phosphate-buffered saline (PBS; Ricca Chemical, R58190001A) at 50 mL/min, followed by perfusion of 2 L of 4% formalin, prepared by diluting 10% neutral buffered formalin (Fisher, 22-110-875) with phosphate-buffered saline, at the same rate of 50 mL/min. The head was then removed and submerged in a separate stock of 4% formalin for at least 24 hours to ensure adequate fixation.

After fixation, the brain was extracted by removing the overlying bone and connective tissue using a motorized band saw. The brain was cryoprotected with 30% sucrose in PBS and frozen in isopentane cooled with dry ice. Tissue was sectioned coronally at 50 µm using an American Optical 860 sliding freezing microtome, and free-floating sections were collected for subsequent staining procedures.

#### Hematoxylin and eosin (H&E) staining

Tissue sections were mounted on gelatin-coated slides and allowed to dry overnight. Slides were rinsed with distilled water and immersed in Gill II hematoxylin (Sigma, GHS216) for ∼2 min, followed by three washes in tap water. To enhance nuclear staining, slides were blued in Scott’s Tap Water Substitute (∼10 dips, 30 s; Newcomer Supply, 1380C), then washed three additional times in tap water. Slides were immersed in 70% ethanol (Fisher, 01-338-169) and stained with modified eosin (Abcam, ab246824) for ∼3 min. After rinsing in distilled water, slides were dehydrated in two changes of 95% ethanol (1 min each), followed by two changes of 100% ethanol (10 dips each). Clearing was performed in three changes of xylene (10 dips each; Fisher, X3P1GAL) and coverslips were mounted using DPX medium (Sigma, 06522).

#### Cresyl violet staining

Tissue sections were mounted on gelatin-coated slides and allowed to dry overnight. Slides were rinsed with distilled water and dehydrated through a graded ethanol series (50%, 70%, 90%, and 100%, 5 min each; Fisher, 01-338-169). Lipids were removed by immersing the slides in xylene (Fisher, X3P1GAL) for 5 min. Slides were then rehydrated through a reverse ethanol series (100%, 90%, 70%, 50%, and distilled water, 5 min each). Staining was performed using 0.5% cresyl violet acetate solution (Sigma, C5042) for ∼3 min. Excess dye was removed by a brief rinse in distilled water. Differentiation was carried out in a graded ethanol series: 70%, 90%, and 100% ethanol for ∼2 min each (times adjusted as needed for optimal contrast). Slides were cleared in two changes of xylene (5 min each) and coverslips were mounted using DPX medium (Sigma, 06522).

#### Imaging and digitization

Slides were imaged at high power using a Nikon Eclipse 80i microscope equipped with 2–20× Plan Apo objectives. Low-power images were taken with a Leitz M5A stereomicroscope at 2–3× magnification. Images were captured using a CCD camera (Lumenera Infinity 3S-UPM) and digitized using Metamorph software (Molecular Devices). Brightness, contrast, and color were adjusted linearly using Adobe Photoshop (Creative Cloud 2025).

## RESULTS

### Infarct characterization by T1-weighted MRI and histology

Longitudinal structural MRI and postmortem histology confirmed the presence of focal lesions within the left internal capsule following ET-1 injection (Figure 3). Pre-infarct T1-weighted MRI scans showed no abnormalities at or near the planned injection site (Figures 3A, S3A, S4A; left column). At 1–3 weeks post-infarct, T1-weighted images exhibited hypointense signal changes near the ET-1 injection centroid (Figure 3A, S3A, S4A; middle columns). By 7 weeks post-infarct, the region appeared hyperintense relative to surrounding white matter, consistent with chronic infarction (Aoe et al., 2006; Weston et al., 2022) (Figure 3A, S3A, S4A; right column). Hypointense and hyperintense changes were evident in both coronal and axial slices aligned to consistent anatomical coordinates.

Histological examination of the left hemisphere confirmed the presence of a well-demarcated lesion restricted to white matter tracts of the internal capsule in Mk-A (Figure 3B). The infarct core was characterized by pallor, vacuolization, loss of normal white matter architecture, reduced cellularity, and the presence of fragmented eosinophilic debris (Figure 3C), features consistent with ischemic tissue damage. In the peri-infarct region, increased cellular density was observed, suggestive of reactive gliosis (Figure 3D). Cresyl violet staining further demonstrated cellular loss and pallor within the infarct core (Figure 3E).

In Mk-D, structural MRI revealed some lesion extension into the adjacent putamen, although the injection centroid remained centered on the internal capsule (Figure S3A). Mk-F showed a more restricted lesion pattern similar to Mk-A, with signal changes mostly localized to the internal capsule (Figure S4A). Taken together, structural MRI and histological findings indicate that infarcts were centered on the targeted white matter region of the internal capsule, though lesion spread into adjacent subcortical gray matter was observed in one subject.

### Proximal forelimb impairment post-infarct

Performance on the center-out joystick task was used to evaluate effects of internal capsule infarct on proximal forelimb control. Trial attempt rates showed minimal raw differences between pre- and post-infarct periods for all animals (Figure 4B for Mk-A, Figure S1A for Mk-D, Figure S2A for Mk-F). After fitting generalized linear mixed models (GLMMs) accounting for per-session variability, predicted probabilities of trial attempts did not differ significantly between pre- and post-infarct periods for Mk-A (*p* = 0.29), Mk-D (*p* = 0.10), or Mk-F (*p* = 0.27). These results suggest that motivation to perform the task was not substantially affected by infarct, as animals remained equally likely to attempt trials between pre- and post-infarct periods.

**Figure 4.**
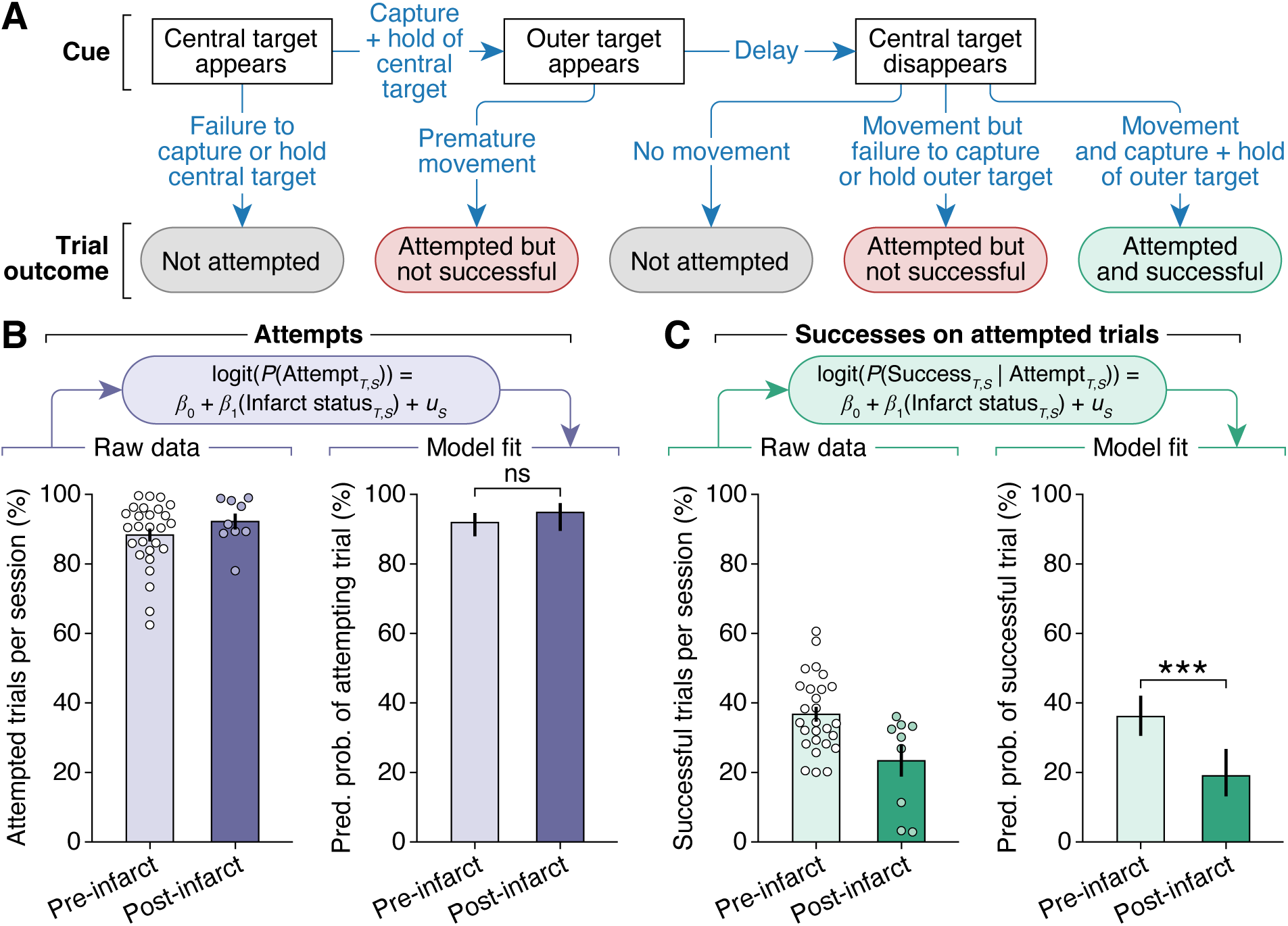
Center-out task performance for an exemplar animal (Mk-A). **A.** Flowchart illustrating the sequence of trial events and possible outcomes in the proximal forelimb center-out joystick task. Task trials are categorized as either not attempted, attempted but not successful, or attempted and successful. **B.** Raw differences in attempt percentage between pre- and post-infarct trials are minimal (left; mean ± SEM). After fitting the data to a generalized linear mixed model (GLMM) that accounts for per-session random effects, predicted probabilities of trial attempts are not significantly different between pre-and post-infarct periods (*p* = 0.29; right; estimated marginal means with 95% confidence intervals). **C.** When considering only attempted trials, raw success percentages appear to be reduced between post-infarct compared to pre-infarct (left; mean ± SEM). After fitting the data to a GLMM that accounts for per-session random effects, predicted probabilities of success for attempted trials differ significantly between pre- and post-infarct periods (*p* = 0.0007; right; estimated marginal means with 95% confidence intervals). Data in **B–C** are from a single animal (Mk-A). Statistical significance for all panels: not significant (ns), *p* < 0.001 (***).

In contrast, success rates among attempted trials were reduced post-infarct for some animals. For Mk-A, predicted probabilities of success were significantly lower following infarct (*p* = 0.0007; Figure 4C), and a significant reduction was also observed for Mk-D (*p* = 0.0095; Figure S1B). In Mk-F, success rates showed no significant difference between pre- and post-infarct periods (*p* = 0.18; Figure S2B). These findings indicate that, although animals continued to attempt trials, the ability to successfully complete the movement was impaired in two of the three subjects (Mk-A and Mk-D).

Joystick kinematics were further analyzed to characterize cursor trajectories on successful trials. Trajectory path lengths were significantly increased post-infarct in Mk-A (*p* = 0.0015; Figure 5B), whereas no significant differences were observed for Mk-D (*p* = 0.30; Figure S1C) or Mk-F (*p* = 0.29; Figure S2C). For Mk-A, this result was further supported by an observed widening of the 95% prediction regions around cursor paths early post-infarct, followed by reduction in variability by the late post-infarct period (Figure 5D). Movement durations, defined as the time between departure from the central target and capture of the outer target, did not differ significantly between pre- and post-infarct periods in Mk-A (*p* = 0.065; Figure 5C) or Mk-D (*p* = 0.21; Figure S1D). In Mk-F, a slight but statistically significant increase in movement duration was observed post-infarct (*p* = 0.033; Figure S2D).

**Figure 5.**
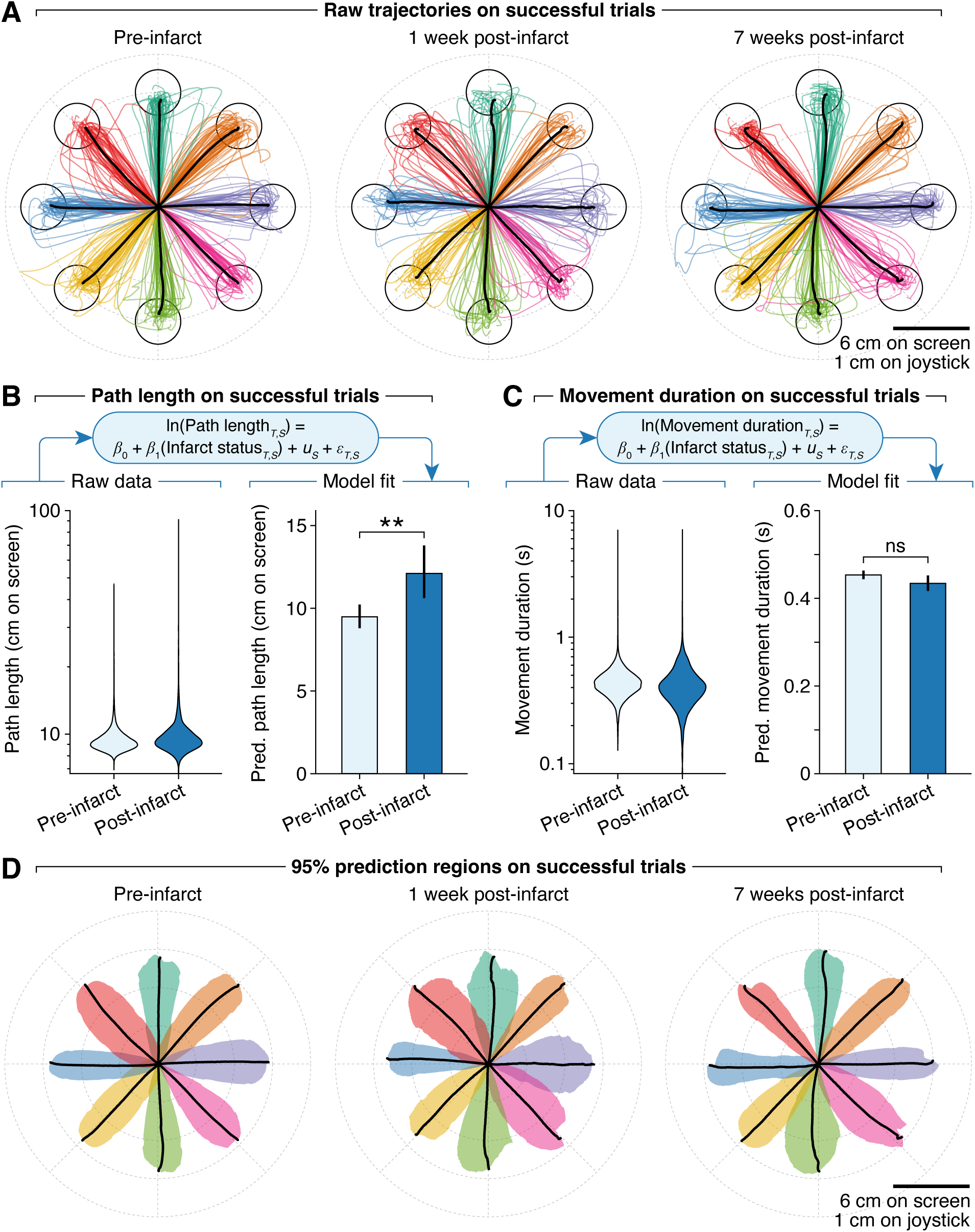
Joystick kinematics for an exemplar animal (Mk-A). **A.** Representative two-dimensional cursor trajectories during the pre-infarct period, 1 week post-infarct, and 7 weeks post-infarct. Trajectories appear tighter and more controlled during pre-infarct and 7-week post-infarct periods, and less controlled at 1 week post-infarct. **B.** Trajectory path length, defined as the cursor distance traveled between the central and outer targets on successful trials. Raw path lengths (left) are analyzed using a GLMM accounting for per-session random effects; predicted path length differs significantly (*p* = 0.0015) between pre- and post-infarct periods (right; estimated marginal means with 95% confidence intervals). **C.** Movement duration, defined as the time taken to move the cursor between the central and outer targets on successful trials. Raw movement durations (left) are similarly analyzed with a GLMM accounting for per-session random effects; predicted movement duration is not significantly different (*p* = 0.065) between pre- and post-infarct periods (right; estimated marginal means with 95% confidence intervals). **D.** Cursor trajectory variability, quantified as the 95% prediction region around trajectories for successful trials, indicating the area within which 95% of future cursor paths would be expected to fall. Prediction regions are consistently sized and shaped during the pre-infarct period, but are irregularly sized and shaped at 1 week and 7 weeks post-infarct. Data in all panels are from a single animal (Mk-A). Statistical significance for all panels: not significant (ns), *p* < 0.01 (**).

Overall, these results suggest that while motivation to engage with the center-out task remained largely intact across animals, impairments in motor function emerged post-infarct. However, the extent and pattern of impairments varied across animals, indicating that the infarcts had differential effects on proximal forelimb control.

### Distal forelimb impairment post-infarct

Performance on the Klüver board task was used to evaluate effects of internal capsule infarct on distal forelimb dexterity and grasping ability. Prior to infarct, animals preferentially used their right (contralesional) hand during trials in which hand choice was unrestricted. Following infarct, all animals showed a significant reduction in right-hand use (Mk-A: *p* = 0.014, Mk-D: *p* < 0.0001, Mk-F: *p* < 0.0001; Figure 6A). Given these shifts in hand preference, subsequent analyses were restricted to trials performed with the right hand to directly assess impairments related to the left-sided infarcts. Pre-infarct task performance showed a consistent difficulty hierarchy across wells, with retrieval time and in-well digit flexions both ranking SW wells as easiest, followed by SN, DW and DN wells (Figure 6B). Because relative performance trends across wells were consistent, subsequent analyses pooled across all well types to examine overall changes in motor function.

**Figure 6.**
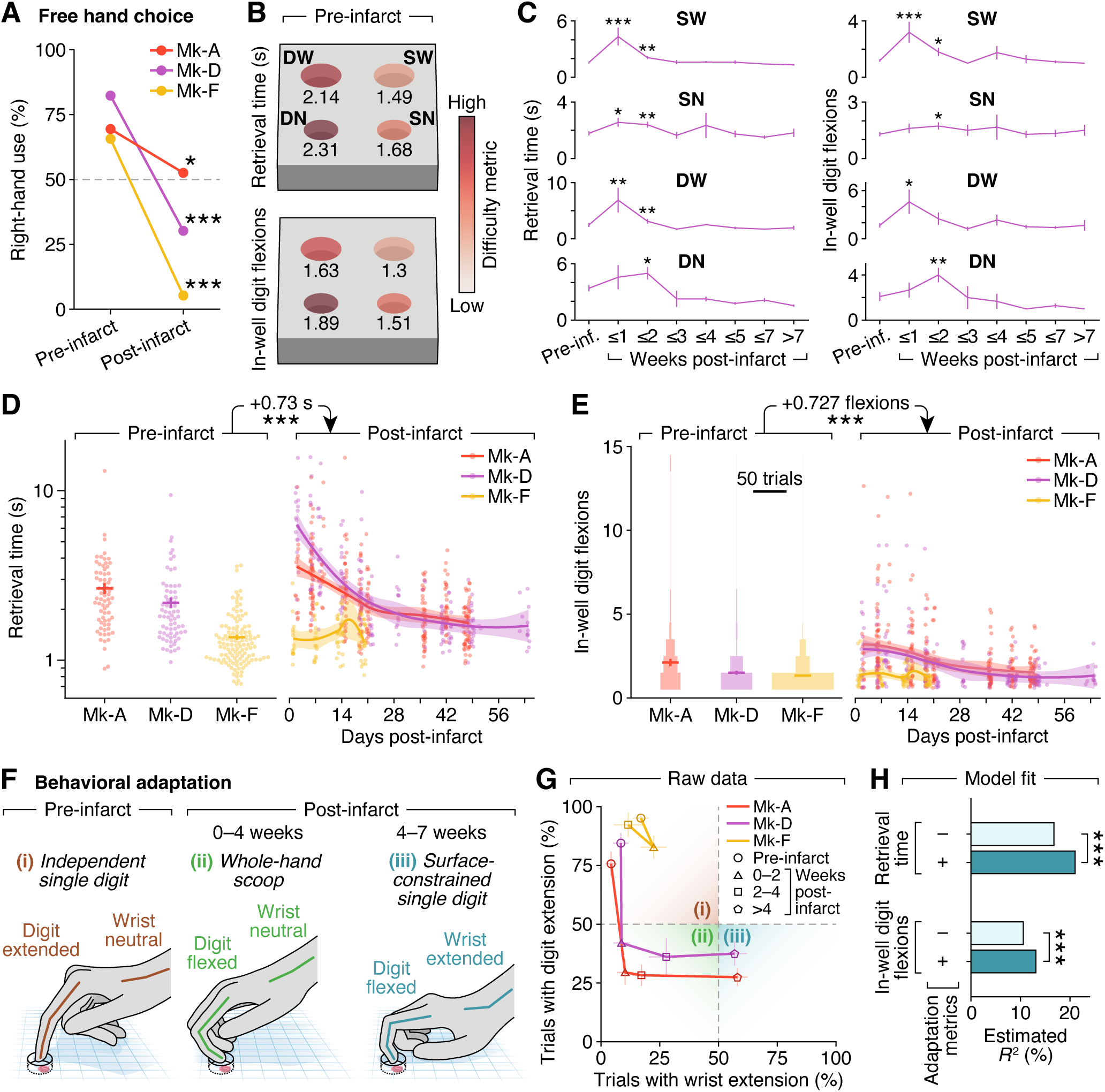
Klüver board task performance. **A.** On trials with unrestricted hand choice, all animals (Mk-A, Mk-D, Mk-F) significantly reduced right-hand use after left internal capsule infarct (Fisher’s exact test; Mk-A: *p* = 0.014, Mk-D: *p* < 0.0001, Mk-F: *p* < 0.0001). Gray dashed line (50%) indicates no hand preference. Subsequent analyses in this figure are limited to right-hand trials only to assess effects of the left-sided infarct. **B.** Heatmaps of pre-infarct mean retrieval time (seconds from movement onset to food contacting the mouth) and mean number of in-well digit flexions reveal a consistent difficulty hierarchy among wells: shallow wide (SW) < shallow narrow (SN) < deep wide (DW) < deep narrow (DN). **C.** Longitudinal retrieval time (left) and digit flexions (right) for each well in a single animal (Mk-D). Motor performance worsened in early post-infarct weeks, then returned to baseline by weeks 3–4. Because performance trends were similar, subsequent analyses pool across all wells. Wilcoxon rank-sum tests (one-sided vs. pre-infarct; *p*-values unadjusted for multiple comparisons) are unstarred if not significant. Data are shown as mean ± SEM. **D–E.** Retrieval time (**D**) and in-well digit flexions (**E**) from pre-infarct trials (left; mean ± SEM) and post-infarct trials (right; smoothed trends with 95% confidence intervals), shown per animal. Mk-A and Mk-D showed early post-infarct impairments with gradual improvement; Mk-F showed minimal change. GLMMs estimate post-infarct increases of 0.73 s in retrieval time (*p* < 0.0001) and 0.727 in digit flexions (*p* < 0.0001), controlling for well type, and days post-infarct by animal. **F–G.** Behavioral adaptation partially explains functional recovery. (i) Pre-infarct: animals used an independent single digit strategy, enabling precise digit flexion. (ii) Early post-infarct (0–4 weeks): animals relied on a whole-hand scoop strategy; impaired digit extension resulted in a flexed hand posture above the Klüver board, limiting digit entry. (iii) Late post-infarct (4–7 weeks): animals adopted a surface-constrained single digit strategy, pressing other digits against the board to improve digit entry and flexion. This adaptation was marked by wrist extension rather than a neutral posture. Illustrations in **F** are representative of postures observed in Mk-A and Mk-D. Data in **G** are shown as proportions ± SE. **H.** Adding wrist and digit extension to GLMMs significantly improves model fit for both retrieval time and digit flexions (likelihood-ratio tests, *p* < 0.0001), indicating that behavioral adaptation explains additional variance in task performance. Statistical significance for all panels: *p* < 0.05 (*), *p* < 0.01 (**), *p* < 0.001 (***).

Motor impairments in distal dexterity were evident early after infarct. Retrieval times, defined as the duration from movement onset to food contacting the mouth, and in-well digit flexions, defined as the number of flexion movements performed while the digits were inside the well, both increased during early post-infarct sessions (Figure 6C). GLMMs revealed significant post-infarct increases in retrieval time (estimate = 0.73 s, *p* < 0.0001) and digit flexions (estimate = 0.727 flexions, *p* < 0.0001) after controlling for well type and days post-infarct by animal (Figures 6D–E). These patterns were observed in both Mk-A and Mk-D, whereas Mk-F showed minimal post-infarct change in either metric.

Video observation of food retrieval strategies indicated that animals changed their motor behavior post-infarct. Pre-infarct, animals predominantly used an independent single-digit strategy, characterized by precise, isolated digit-II flexion within the wells (Figure 6F, left). In the early post-infarct period (0–4 weeks), animals frequently exhibited a flexed hand posture above the Klüver board, impairing the ability to extend digit II into the target well; movements during this period were instead characterized by a whole-hand scooping strategy (Figure 6F, middle). By the late post-infarct period (4–7 weeks), animals adapted by employing a surface-constrained single-digit strategy, characterized by wrist extension and by pressing the flexed digits III–V against the Klüver board surface. This posture provided hand stabilization and allowed digit II to enter the wells at a more board-orthogonal angle, facilitating improved distal interphalangeal (DIP) joint flexion during retrieval (Figure 6F, right). Although this adaptation was not observed on every trial, it occurred in more than 50% of right-hand retrievals in both Mk-A and Mk-D (Figure 6G). Trial-wise GLMMs incorporating the two identified adaptation metrics (wrist extension and digit-II extension) significantly improved model fits for both retrieval time and digit flexions, as assessed by likelihood-ratio tests (*p* < 0.0001 for each; Figure 6H). For retrieval time, the estimated 𝑅^2^increased from 16.7% without adaptation metrics to 21.0% with adaptation metrics; for digit flexions, the estimated 𝑅^2^increased from 10.6% without adaptation metrics to 13.1%. These findings suggest that behavioral adaptations contributed to improvements in task performance during the late post-infarct period.

Together, these results show that distal forelimb control was impaired following internal capsule infarct, and that functional recovery over time was associated with the emergence of alternative task strategies that improved digit flexion and food retrieval.

### Infarct characterization by T2-weighted MRI and DTI

T2-weighted MRI scans showed signal changes consistent with infarct evolution described in T1-weighted imaging. Pre-infarct T2 signal near the planned injection site was normal. At 1–3 weeks post-infarct, T2 signal became hyperintense, consistent with acute ischemic changes, and by 7 weeks, the region became hypointense, consistent with chronic infarction (Figure 7A). To estimate infarct volume at the 1-week post-infarct timepoint, hyperintense voxels exceeding an animal-specific intensity threshold were counted and multiplied by voxel volume (0.125 mm^3^). These estimates are reported in Table 5. Mk-D exhibited the largest infarct volume, followed by Mk-A, then Mk-F. This ordering corresponded with the severity of post-infarct motor impairments observed on the Klüver board task: Mk-D showed the most pronounced increase in retrieval time and digit flexions, followed by Mk-A, while Mk-F showed only mild post-infarct changes (Figure 6D–E).

**Figure 7.**
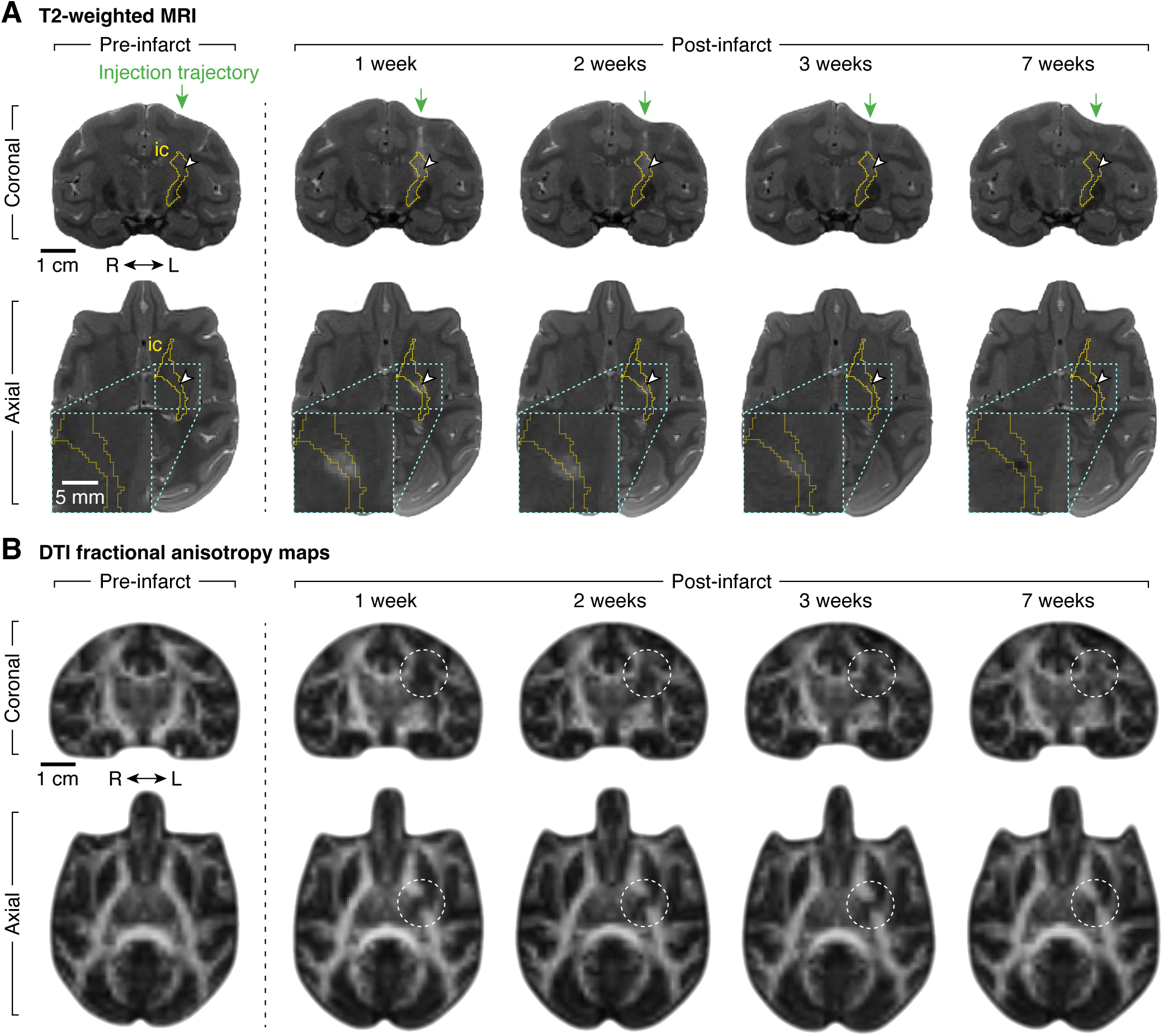
T2-weighted MRI and diffusion tensor imaging (DTI) characterization of ET-1-induced internal capsule infarct in an exemplar animal (Mk-A). **A.** Longitudinal T2-weighted MRI scans illustrating infarct evolution in the internal capsule. Each column shows coronal (top row) and axial (bottom row) slices at consistent atlas-aligned coordinates. Internal capsule (ic) is outlined in yellow; ET-1 injection centroid is marked by a white arrowhead (planned in pre-infarct, actual in post-infarct images). Coronal slices include a green arrow denoting the vertical injection trajectory. Axial slices include a bottom-left inset enlarging the region near the injection site. Anatomical orientation marker indicates right (R) and left (L) hemispheres. Pre-infarct scans exhibit normal T2 signal intensity. Post-infarct scans (1–3 weeks) show transient hyperintensity near injection centroid, consistent with edema and acute ischemic changes. By 7 weeks, the region appears hypointense, indicative of chronic infarction and tissue loss. **B.** DTI fractional anisotropy (FA) maps at the same coordinates and time points as in **A**. Bright voxels represent intact, directionally coherent white matter fibers; dark voxels represent disrupted fiber integrity. Dashed white circles highlight infarct-associated white matter disruption. Pre-infarct scan shows normal FA within the internal capsule. Post-infarct scans (1–3 weeks) demonstrate marked reductions in FA near the ET-1 injection centroid, consistent with acute ischemic alterations to white matter. By 7 weeks, FA partially recovers. Images in all panels are from a single animal (Mk-A).

Additional structural characterization was provided by DTI. Pre-infarct fractional anisotropy (FA) maps showed bright anisotropic signal within the internal capsule, consistent with intact white matter fiber organization (Figure 7B, left column). At 1–3 weeks post-infarct, marked reductions in FA were observed near the injection centroid, consistent with alterations to white matter (Figure 7B, middle columns). By 7 weeks post-infarct, partial recovery of FA signal was observed, although values remained lower than pre-infarct levels (Figure 7B, right column).

## DISCUSSION

The primary findings of this study are fivefold. First, we established a reproducible white matter stroke model in Rhesus macaques by targeting the internal capsule with ET-1 injections. Infarcts were confirmed through longitudinal MRI and histology to be spatially localized to the intended subcortical region, with preservation of overlying gray matter. Second, animals exhibited variable impairments in proximal forelimb function following infarct, as measured by a joystick-based center-out task. While motivation to engage with the task remained intact, some animals showed reduced success rates and increased path lengths and movement durations on successful trials, indicating mild to moderate proximal motor deficits. Third, more robust and consistent impairments were observed in distal forelimb dexterity using a Klüver board task. All animals showed a significant reduction in use of the contralesional hand, and difficulty metrics such as retrieval time and in-well digit flexions increased following infarct in two animals, reflecting loss of fine motor control. Fourth, animals demonstrated behavioral adaptation over time, specifically through a strategy in which flexed digits were stabilized against the Klüver board surface by extending the wrist, enabling improved food retrieval. Incorporating these adaptation metrics in statistical models of motor function significantly increased explanatory power. Fifth, infarct volumes varied across animals, and larger infarcts were associated with greater impairment in motor task performance.

This NHP stroke model is both novel and clinically relevant for several reasons. In humans, loss of distal hand and finger function is among the most debilitating consequences of motor stroke, with high impact on independence and quality of life (Houwink et al., 2013; Lang et al., 2013). Fine motor control of the fingers is largely mediated by CST fibers originating from the M1 hand area, which project monosynaptically to spinal motor neurons innervating distal musculature (Strick et al., 2021; Lemon, 2004; Bortoff & Strick, 1993; Lawrence & Kuypers, 1968). Importantly, these CST projections are densely packed within subcortical white matter tracts, including the posterior limb of the internal capsule (Morecraft et al., 2002). White matter infarcts in these regions are often associated with worse motor outcomes and slower recovery trends than cortical infarcts (Wang et al., 2016; Corbetta et al., 2015; Schiemanck et al., 2008). Because the mechanisms underlying white matter stroke remain poorly understood, yet are associated with disproportionately severe disability, a model focused on white matter injury can help clarify its specific role in post-stroke motor deficits. Furthermore, the majority of clinical strokes involve subcortical areas (Wang et al., 2016; Corbetta et al., 2015), highlighting the translational relevance of targeting these regions.

In the macaque brain, the internal capsule can be reliably targeted using subject-specific structural MRI, enabling reproducible subcortical lesioning while preserving surrounding gray matter. Compared to cortical lesions, white matter infarcts as implemented in this study also leave intact cortical structures for concurrent monitoring or therapeutic modulation, making this model compatible with multimodal approaches such as electrocorticography or neurostimulation. Additionally, NHPs offer critical advantages over other animal models due to their anatomical and functional homology to humans, including differentiation of premotor areas and the presence of direct monosynaptic CST projections (Nudo & Masterton, 1990). Monosynaptic projections to the ventral horn are found in only a subset of NHP species, including Rhesus macaques, and are thought to underlie the capacity for dexterous finger movements (Bortoff & Strick, 1993). For these reasons, the Stroke Therapy Academic Industry Roundtable (STAIR) committee has recommended the use of NHPs for preclinical stroke studies to improve translational fidelity (Fisher et al., 2009). Our model leverages these species-specific features to selectively disrupt white matter fibers within the internal capsule, providing a platform for probing mechanisms of motor impairment, adaptation, and recovery with clear relevance to human stroke.

This model also recapitulates a key behavioral phenomenon observed in human stroke recovery: a shift in hand preference away from the more-affected limb for tasks requiring fine motor control (Waddell et al., 2017). All three macaques demonstrated reduced use of the contralesional (right) hand on the Klüver board task following left internal capsule infarct, consistent with disruption of M1-originating CST output pathways. Deficits in retrieval time and in-well digit flexions further indicated impaired distal motor control. Importantly, standard performance metrics such as retrieval time alone were insufficient to fully characterize post-infarct changes; detailed observation of movement patterns was necessary to reveal the evolving strategies animals used during functional recovery. While the Klüver board task was highly sensitive to these impairments, the center-out joystick task yielded more variable results. Although designed to assess proximal reaching, successful performance on the center-out task also required distal grasp and stabilization, engaging multiple motor components across the forelimb. Post-infarct outcomes on this task varied across animals, with some showing clear impairments and others exhibiting less pronounced change. This variability reinforces the importance of selecting behavioral tasks that are both anatomically specific and sensitive to inter-subject differences in motor impairment.

Variability in motor outcomes across animals was likely shaped by a combination of anatomical, behavioral, and infarct-related factors. One of the three animals (Mk-F) exhibited notably milder impairments despite receiving a targeted infarct to the internal capsule. These differences may reflect individual variation in white matter anatomy, particularly the spatial organization of descending motor fibers and their projection targets. Differences in post-injection diffusion kinetics may also have influenced infarct distribution and the degree of disruption to motor pathways. In addition, this animal displayed consistently higher levels of baseline motor activity in its home environment, with faster, more frequent movements and heightened responsiveness to external stimuli. These behavioral tendencies may have promoted greater spontaneous use of the affected limb following infarct, contributing to functional recovery through increased non-task-related motor activity.

Variability in infarct size and distribution also appeared to track with differences in functional outcomes, with more extensive infarcts associated with more pronounced motor impairment. Despite careful targeting, some infarct extension into the basal ganglia was observed on early post-infarct structural MRI in one animal (Mk-D). This variability in the pattern of infarct spread across animals is consistent with human subcortical stroke, where ischemic injury involving the internal capsule typically results from occlusion of lenticulostriate arteries and exhibits variable involvement of neighboring basal ganglia structures (Marinković et al., 2001; Decavel et al., 2012). Thus, while inter-subject differences were present, the infarct patterns observed in this study are broadly coincident with clinically relevant subcortical stroke presentations.

Further supporting the translational relevance of this model, a characteristic flexed posture of the hand and fingers was observed in our animals following internal capsule infarct (Figure 6F, middle), mirroring the flexor synergy pattern frequently seen in human patients with subcortical stroke (O’Sullivan et al., 2019; Sawner & LaVigne, 1992). This posture is commonly attributed to selective disruption of CST output and relative preservation of descending extrapyramidal pathways, including the reticulospinal, rubrospinal, and vestibulospinal tracts, which are thought to favor flexor tone in the upper limb (Lawrence & Kuypers, 1968; Riddle et al., 2009; Baker, 2011; Olivares-Moreno et al., 2021). Given that wrist extensors are extrinsic to the hand and may be represented in a portion of the internal capsule distinct from the area most affected by the infarct (Casteleyn & Bakker, 2021; Morecraft et al., 2002), it is plausible that the motor units controlling these muscles were relatively spared and remained available for task execution. Notably, wrist extension used to orient and stabilize the distal forelimb is consistent with locomotor hand postures described in palmigrade-capable primates, including macaques, during ethologically relevant behaviors such as foraging and climbing (Orr, 2017). The observed behavioral adaptation may therefore reflect increased reliance on pre-existing motor routines that were co-opted to accomplish the Klüver board task in the absence of the preferred retrieval strategy. These preserved motor routines may have been mediated in part by efferent extrapyramidal pathways, particularly the reticulospinal and rubrospinal tracts, which are involved in goal-directed forelimb movements and can compensate for CST disruption (Riddle et al., 2009; Baker, 2011; Olivares-Moreno et al., 2021). Histological evidence of partial sparing of M1-originating fibers is consistent with the reuse of such routines, though it does not establish which motor pathways supported the observed behavior. Nevertheless, the emergence of an adaptive motor strategy in response to impaired task-relevant motor control mirrors clinical stroke phenotypes and suggests that this model may capture behaviorally meaningful features of post-infarct motor function.

The emergence of behavioral adaptation in this model is reminiscent of a broader clinical phenomenon: human stroke patients frequently adopt alternative movement strategies to regain functional performance, rather than restoring the original motor patterns used before injury (Roby-Brami et al., 2021). For example, constraint-induced movement therapy is based on restricting compensatory movements to promote use of the impaired limb that more closely reflects pre-stroke motor control (Kwakkel et al., 2015). Likewise, some robotic rehabilitation approaches aim to detect and reduce compensatory movements, guiding patients toward pre-stroke-like movement patterns (Zhi et al., 2018). The distinction between functional restoration and compensation carries practical clinical consequences, as persistent use of maladaptive movements can contribute to joint stress, overuse injuries, and poorer long-term recovery outcomes (Takeuchi & Izumi, 2012).

These considerations point to important directions for future work. First, stimulation-based mapping approaches could be used to refine infarct targeting, allowing more precise disruption of motor representations relevant to specific task components. Second, future investigations may benefit from assessing a wide panel of naturalistic behaviors that reflect ethologically relevant motor routines, as these contexts could more readily reveal adaptive strategies that emerge following focal white matter infarct. Notably, in the present study, changes in hand preference post-infarct did not appear to be directly related to motor capacity in the affected limb, reflecting the broader observation that lab-based performance metrics may not fully capture behavior in more naturalistic settings (Hayward et al., 2016). Studying these behaviors in greater detail may enable more precise characterization of how animals modify their movements post-infarct using preserved motor pathways. Selecting tasks that are more likely to generalize across environments may further enhance the clinical relevance of this model, since meaningful recovery for patients is ultimately defined by improved performance in activities of daily living rather than success on narrowly defined behavioral tests.

Several limitations of this study should be acknowledged. First, infarct volumes in our model were smaller than those typically observed in clinical cases of subcortical stroke with white matter involvement. Although the lesions were well localized and reproducible, the relatively limited infarct size may not fully capture the range or severity of motor deficits seen in human patients. Second, animals underwent an extended training period on both behavioral tasks followed by a prolonged pause in training prior to infarct induction. Although this pause was partially intended to mitigate overtraining, the overall duration of task exposure may still have reduced sensitivity to subtle impairments or enabled compensation through well-established motor routines. Third, all animals were young adults (5–10 years of age), which corresponds to an earlier life stage relative to typical human stroke patients, accounting for species lifespan differences. Human stroke patients are often significantly older and may experience more limited recovery due to age-related factors. These limitations guide appropriate interpretation of our findings, acknowledging that the experimental constraints required to prioritize reproducibility and precise behavioral characterization inherently limit the ability to model the full complexity and variability of human stroke.

## ACKNOWLEDGEMENTS

We thank Donna Reedy for outstanding animal care and handling and Nicholas Luczak for helping port the center-out task to the BCI2000 platform. We are grateful to Catherine E. Lang and Alexandre R. Carter for guiding the analysis of motor function, Cihat Eldeniz and Babatunde Adeyemo for expertise in MRI data acquisition and analysis, April Ratliff and Phillip Demarest for helpful contributions to analyses and figures, and Krikor Dikranian for assistance in interpreting behavioral and histology findings. This work was funded by the following grants: NIH/NINDS R01-NS101013; NIH T32-GR0019882, T32-GR0024755; NIH/NINDS T32-NS126157; NIH/NIBIB P41-EB018783, R01-EB026439; NIH/NINDS U24-NS109103, U01-NS128612.

## AUTHOR CONTRIBUTIONS

DWM, ECL, HB, SAA, and SSS conceptualized and designed the study. DWM, SAA, SSS, DTB, and LB performed surgical procedures, MRI scans, and behavioral experiments. PB provided computational support and assistance with the BCI2000 platform. RLM and AB performed histological procedures. SSS and SAA performed formal data analysis. SSS drafted the manuscript. SAA, DTB, HB, ECL, and DWM revised the manuscript. SSS and SAA prepared the figures. All authors reviewed and approved the final manuscript.

## DISCLOSURES

Eric C. Leuthardt has the following disclosures. Stock ownership: Neurolutions, Face to Face Biometrics, Caeli Vascular, Acera, Sora Neuroscience, Inner Cosmos, Aurenar, Petal Surgical, Inflexion Vascular, Cordance Medical, Silent Surgical. Consultant: Monteris Medical, E15, Neurolutions. Licensing from Intellectual Property: Neurolutions, Caeli Vascular, Inner Cosmos. Licensing/Product Development Agreements or Royalties for inventions/IP: Intellectual Ventures, Sora Neuroscience, Inner Cosmos, Neurolutions, Aurenar.

Daniel W. Moran has the following disclosures. Stock ownership: Neurolutions, Inner Cosmos. Licensing from Intellectual Property: Neurolutions.

David T. Bundy has the following disclosure. Stock ownership: Neurolutions.

Washington University in St. Louis owns equity in Neurolutions.

---

**Figure S1.**
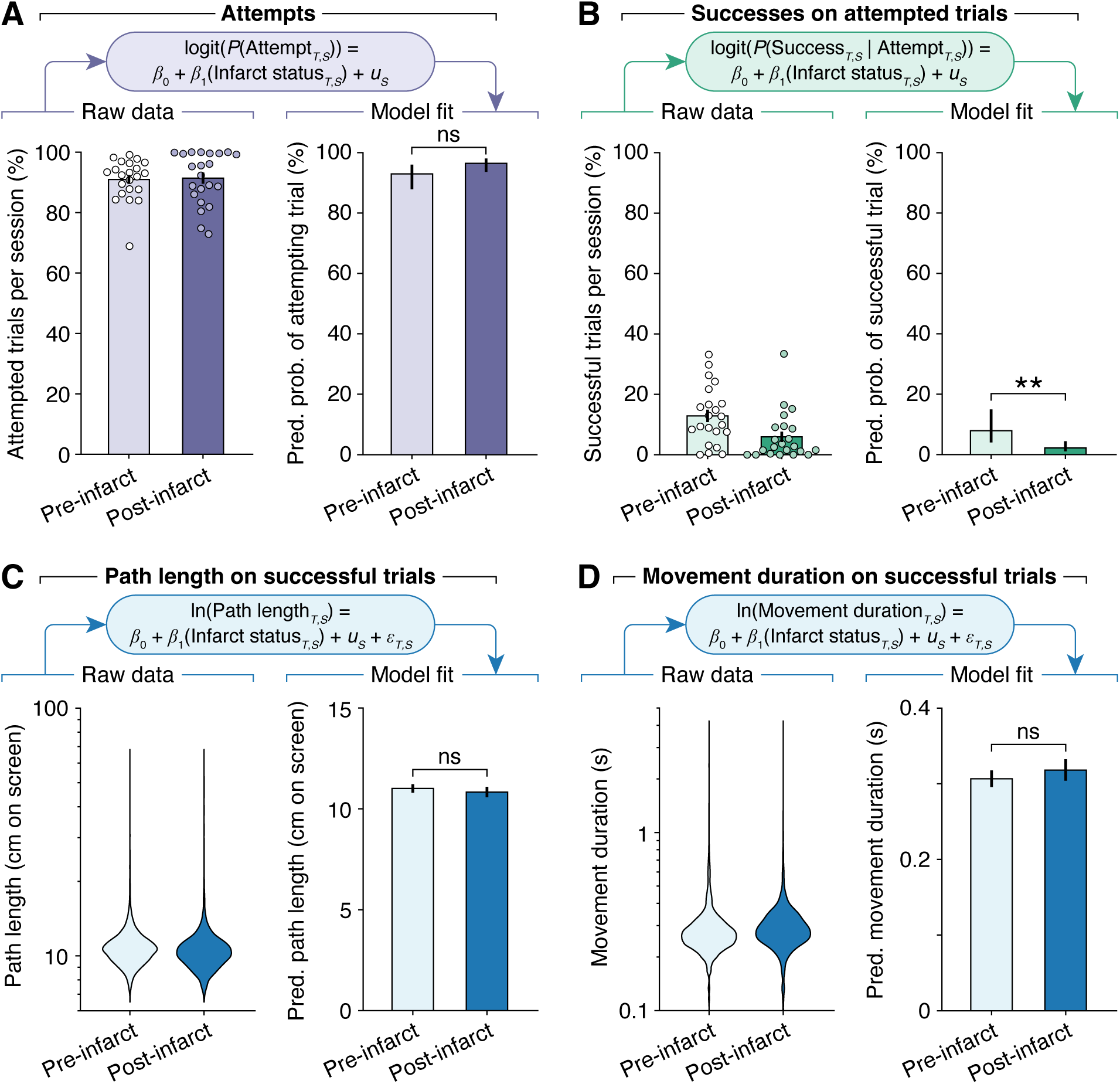
Center-out task performance for Mk-D. **A.** Raw differences in attempt percentage between pre- and post-infarct trials are minimal (left; mean ± SEM). After fitting the data to a generalized linear mixed model (GLMM) that accounts for per-session random effects, predicted probabilities of trial attempts are not significantly different between pre-and post-infarct periods (*p* = 0.10; right; estimated marginal means with 95% confidence intervals). **B.** When considering only attempted trials, raw success percentages appear to be reduced between post-infarct compared to pre-infarct (left; mean ± SEM). After fitting the data to a GLMM that accounts for per-session random effects, predicted probabilities of success for attempted trials differ significantly between pre- and post-infarct periods (*p* = 0.0095; right; estimated marginal means with 95% confidence intervals). **C.** Trajectory path length, defined as the cursor distance traveled between the central and outer targets on successful trials. Raw path lengths (left) are analyzed using a GLMM accounting for per-session random effects; predicted path length does not differ significantly (*p* = 0.30) between pre- and post-infarct periods (right; estimated marginal means with 95% confidence intervals). **D.** Movement duration, defined as the time taken to move the cursor between the central and outer targets on successful trials. Raw movement durations (left) are similarly analyzed with a GLMM accounting for per-session random effects; predicted movement duration is not significantly different (*p* = 0.21) between pre- and post-infarct periods (right; estimated marginal means with 95% confidence intervals). Statistical significance for all panels: not significant (ns), *p* < 0.01 (**).

**Figure S2.**
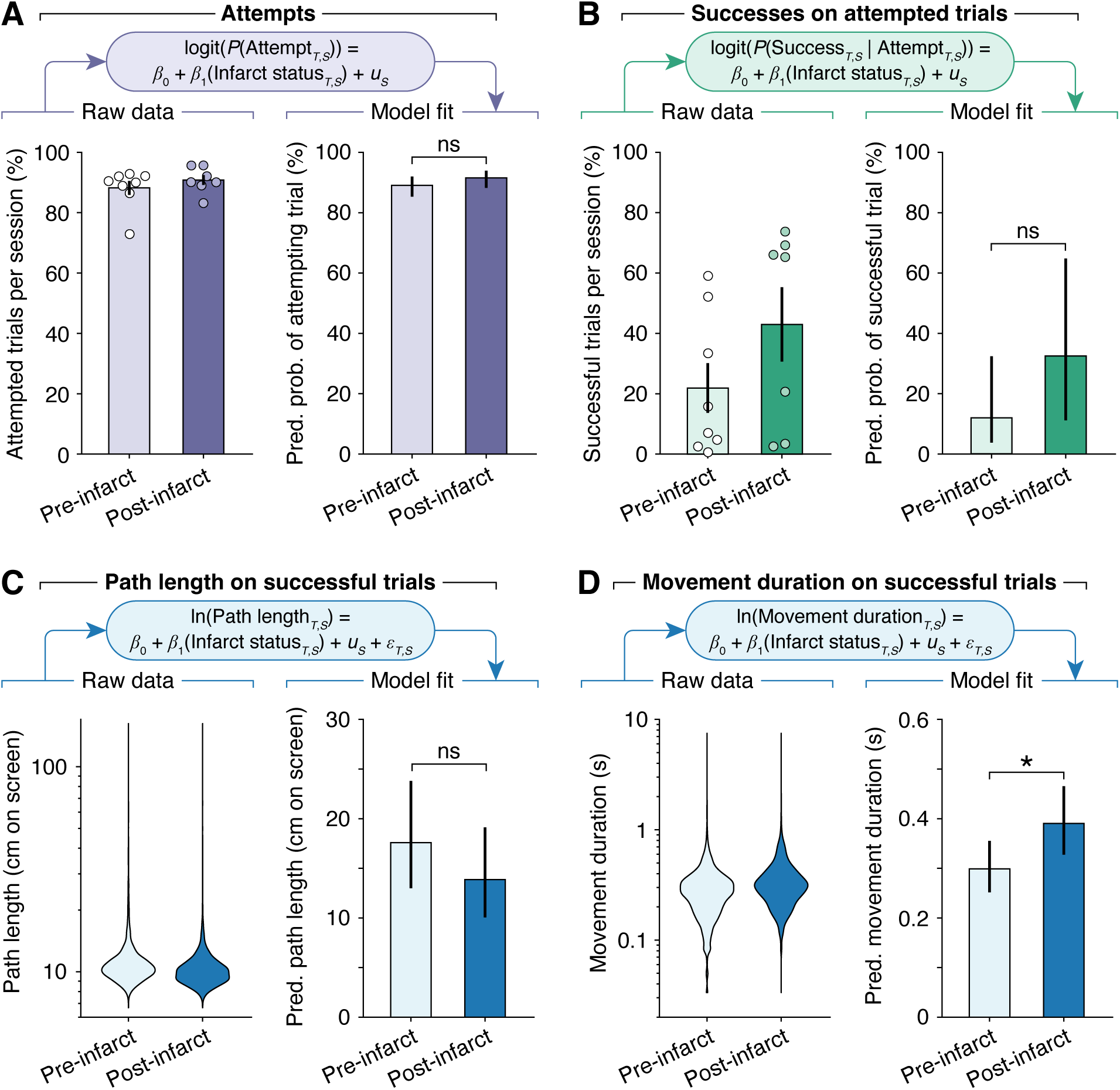
Center-out task performance for Mk-F. **A.** Raw differences in attempt percentage between pre- and post-infarct trials are minimal (left; mean ± SEM). After fitting the data to a generalized linear mixed model (GLMM) that accounts for per-session random effects, predicted probabilities of trial attempts are not significantly different between pre-and post-infarct periods (*p* = 0.27; right; estimated marginal means with 95% confidence intervals). **B.** When considering only attempted trials, raw success percentages have similarly high variability in both pre- and post-infarct periods (left; mean ± SEM). After fitting the data to a GLMM that accounts for per-session random effects, predicted probabilities of success for attempted trials do not differ significantly between pre- and post-infarct periods (*p* = 0.18; right; estimated marginal means with 95% confidence intervals). **C.** Trajectory path length, defined as the cursor distance traveled between the central and outer targets on successful trials. Raw path lengths (left) are analyzed using a GLMM accounting for per-session random effects; predicted path length does not differ significantly (*p* = 0.29) between pre- and post-infarct periods (right; estimated marginal means with 95% confidence intervals). **D.** Movement duration, defined as the time taken to move the cursor between the central and outer targets on successful trials. Raw movement durations (left) are similarly analyzed with a GLMM accounting for per-session random effects; predicted movement duration is slightly increased post-infarct, and this difference is statistically significant (*p* = 0.033; right; estimated marginal means with 95% confidence intervals). Statistical significance for all panels: not significant (ns), *p* < 0.05 (*).

**Figure S3.**
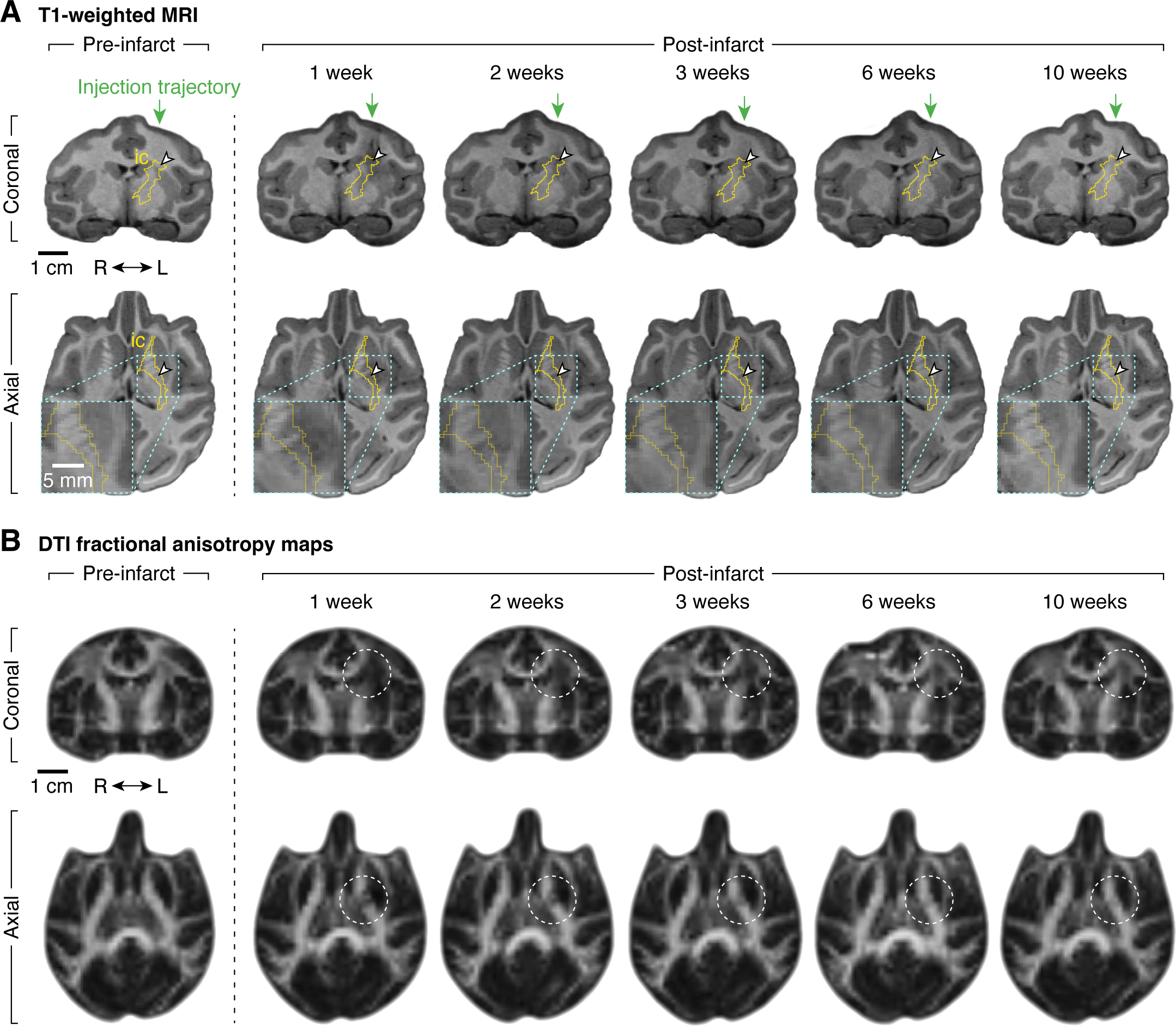
T1-weighted MRI and DTI of Mk-D. **A.** Longitudinal T1-weighted MRI scans illustrating infarct evolution in the internal capsule. Each column shows coronal (top row) and axial (bottom row) slices at consistent atlas-aligned coordinates. Internal capsule (ic) is outlined in yellow; ET-1 injection centroid is marked by a white arrowhead (planned in pre-infarct, actual in post-infarct images). Coronal slices include a green arrow denoting the vertical injection trajectory. Axial slices include a bottom-left inset enlarging the region near the injection site. Anatomical orientation marker indicates right (R) and left (L) hemispheres. Pre-infarct scan shows normal T1 signal near the injection site. Post-infarct scans were acquired at 1, 2, 3, 6, and 10 weeks. Hypointense voxels near the injection centroid are evident at 1–3 weeks and diminish over time; at 6–10 weeks, the region appears slightly hyperintense, consistent with chronic infarct. **B.** DTI fractional anisotropy (FA) maps at the same coordinates and time points as in **A**. Dashed white circles highlight infarct-associated white matter disruption. Pre-infarct scan shows normal FA within the internal capsule. Post-infarct scans (1–3 weeks) show marked reductions in FA near the injection centroid, consistent with acute disruption of white matter integrity. By 10 weeks, FA partially recovers.

**Figure S4.**
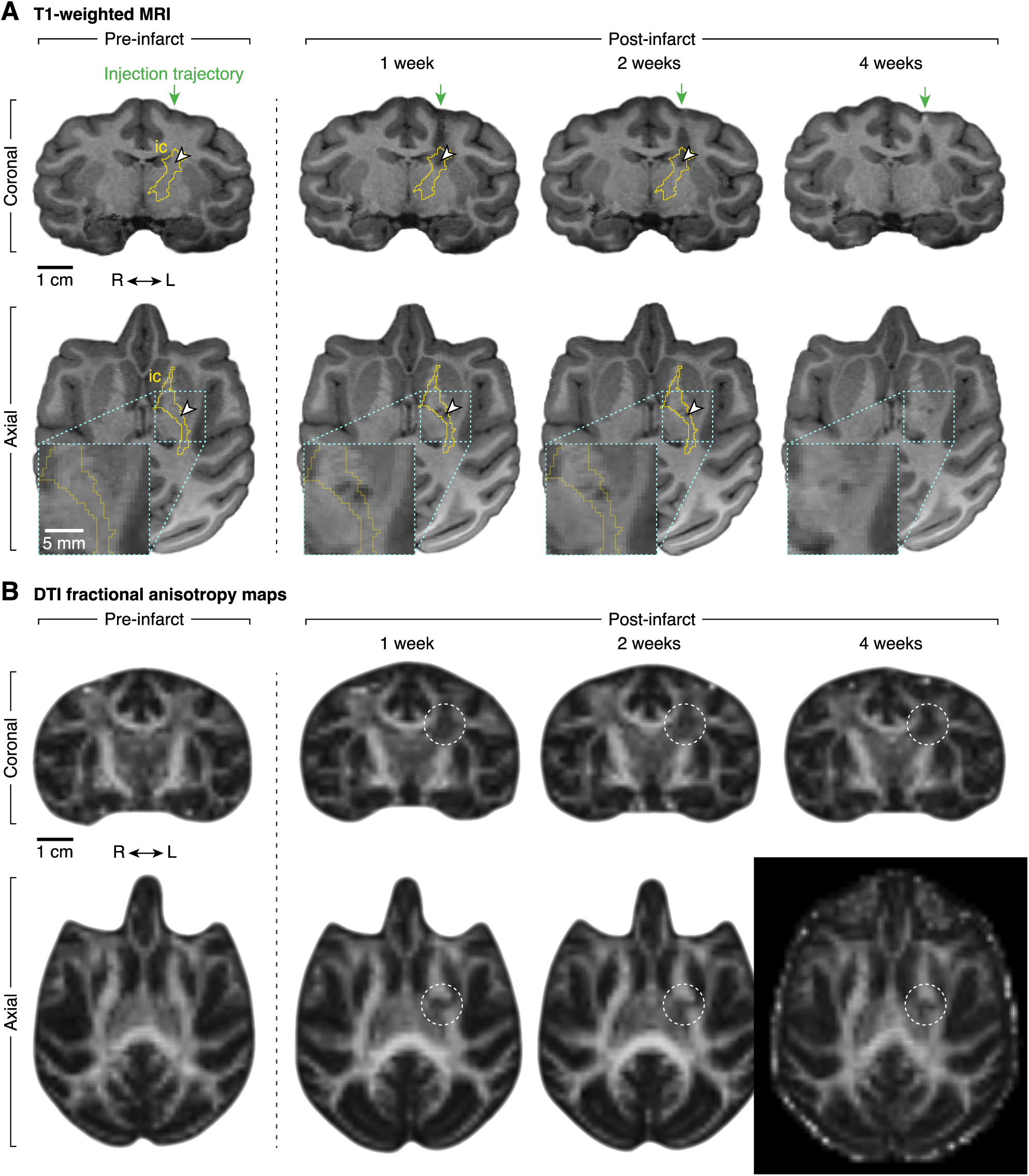
T1-weighted MRI and DTI of Mk-F. **A.** Longitudinal T1-weighted MRI scans illustrating infarct evolution in the internal capsule. Each column shows coronal (top row) and axial (bottom row) slices at consistent atlas-aligned coordinates. Internal capsule (ic) is outlined in yellow; ET-1 injection centroid is marked by a white arrowhead (planned in pre-infarct, actual in post-infarct images). Coronal slices include a green arrow denoting the vertical injection trajectory. Axial slices include a bottom-left inset enlarging the region near the injection site. Anatomical orientation marker indicates right (R) and left (L) hemispheres. Pre-infarct scan shows normal T1 signal near the injection site. Post-infarct scans were acquired at 1, 2, and 4 weeks. Hypointense voxels near the injection centroid are evident at 1–4 weeks and diminish over time. Due to chamber-associated tissue compression, the 4-week scan was not atlas-aligned. **B.** DTI fractional anisotropy (FA) maps at the same coordinates and time points as in **A**. Dashed white circles highlight infarct-associated white matter disruption. Pre-infarct scan shows normal FA within the internal capsule. Post-infarct scans (1–2 weeks) show reductions in FA near the injection centroid, consistent with white matter injury. By 4 weeks, partial recovery of FA is observed.

## REFERENCES

Aoe, H., Takeda, Y., Kawahara, H., Tanaka, A., & Morita, K. (2006). Clinical significance of T1-weighted MR images following transient cerebral ischemia. Journal of the Neurological Sciences, 241(1–2), 19–24. 10.1016/j.jns.2005.10.013

Baker, S. N. (2011). The primate reticulospinal tract, hand function and functional recovery. The Journal of Physiology, 589(23), 5603–5612. 10.1113/jphysiol.2011.215160

Bederson, J. B., Pitts, L. H., Tsuji, M., Nishimura, M. C., Davis, R. L., & Bartkowski, H. (1986). Rat middle cerebral artery occlusion: Evaluation of the model and development of a neurologic examination. Stroke, 17(3), 472–476. 10.1161/01.STR.17.3.472

Bortoff, G. A., & Strick, P. L. (1993). Corticospinal terminations in two new-world primates: Further evidence that corticomotoneuronal connections provide part of the neural substrate for manual dexterity. Journal of Neuroscience, 13(12), 5105–5118. 10.1523/JNEUROSCI.13-12-05105.1993

Carmichael, S. T. (2005). Rodent Models of Focal Stroke: Size, Mechanism, and Purpose. NeuroRX, 2(3), 396–409. 10.1602/neurorx.2.3.396

Carter, A. R., Patel, K. R., Astafiev, S. V., Snyder, A. Z., Rengachary, J., Strube, M. J., Pope, A., Shimony, J. S., Lang, C. E., Shulman, G. L., & Corbetta, M. (2012). Upstream dysfunction of somatomotor functional connectivity after corticospinal damage in stroke. Neurorehabilitation and Neural Repair, 26(1), 7–19. 10.1177/1545968311411054

Casteleyn, C., & Bakker, J. (2021). Anatomy of the Rhesus Monkey (Macaca mulatta): The Essentials for the Biomedical Researcher. In Updates on Veterinary Anatomy and Physiology. IntechOpen. 10.5772/intechopen.99067

Chan, S. S., & Moran, D. W. (2006). Computational model of a primate arm: From hand position to joint angles, joint torques and muscle forces. Journal of Neural Engineering, 3(4), 327–337. 10.1088/1741-2560/3/4/010

Corbetta, M., Ramsey, L., Callejas, A., Baldassarre, A., Hacker, C. D., Siegel, J. S., Astafiev, S. V., Rengachary, J., Zinn, K., Lang, C. E., Connor, L. T., Fucetola, R., Strube, M., Carter, A. R., & Shulman, G. L. (2015). Common Behavioral Clusters and Subcortical Anatomy in Stroke. Neuron, 85(5), 927–941. 10.1016/j.neuron.2015.02.027

Darling, W. G., Pizzimenti, M. A., Rotella, D. L., Peterson, C. R., Hynes, S. M., Ge, J., Solon, K., McNeal, D. W., Stilwell-Morecraft, K. S., & Morecraft, R. J. (2009). Volumetric effects of motor cortex injury on recovery of dexterous movements. Experimental Neurology, 220(1), 90–108. 10.1016/j.expneurol.2009.07.034

Decavel, P., Vuillier, F., & Moulin, T. (2012). Lenticulostriate infarction. Frontiers of Neurology and Neuroscience, 30, 115–119. 10.1159/000333606

DeVetten, G., Coutts, S. B., Hill, M. D., Goyal, M., Eesa, M., O’Brien, B., Demchuk, A. M., & Kirton, A. (2010). Acute Corticospinal Tract Wallerian Degeneration Is Associated With Stroke Outcome. Stroke, 41(4), 751–756. 10.1161/STROKEAHA.109.573287

Edwardson, M. A., Wang, X., Liu, B., Ding, L., Lane, C. J., Park, C., Nelsen, M. A., Jones, T. A., Wolf, S. L., Winstein, C. J., & Dromerick, A. W. (2017). Stroke Lesions in a Large Upper Limb Rehabilitation Trial Cohort Rarely Match Lesions in Common Preclinical Models. Neurorehabilitation and Neural Repair, 31(6), 509–520. 10.1177/1545968316688799

Fisher, M., Feuerstein, G., Howells, D. W., Hurn, P. D., Kent, T. A., Savitz, S. I., & Lo, E. H. (2009). Update of the Stroke Therapy Academic Industry Roundtable Preclinical Recommendations. Stroke, 40(6), 2244–2250. 10.1161/STROKEAHA.108.541128

Friel, K. M., & Nudo, R. J. (1998). Recovery of motor function after focal cortical injury in primates: Compensatory movement patterns used during rehabilitative training. Somatosensory & Motor Research, 15(3), 173–189. 10.1080/08990229870745

Georgopoulos, A. P., Schwartz, A. B., & Kettner, R. E. (1986). Neuronal Population Coding of Movement Direction. Science, 233(4771), 1416–1419. 10.1126/science.3749885

Gladstone, D. J., Black, S. E., & Hakim, A. M. (2002). Toward Wisdom From Failure. Stroke, 33(8), 2123–2136. 10.1161/01.STR.0000025518.34157.51

Hayward, K. S., Eng, J. J., Boyd, L. A., Lakhani, B., Bernhardt, J., & Lang, C. E. (2016). Exploring the Role of Accelerometers in the Measurement of Real World Upper-Limb Use After Stroke. Brain Impairment, 17(1), 16–33. 10.1017/BrImp.2015.21

Herbert, W. J., Powell, K., & Buford, J. A. (2015). Evidence for a role of the reticulospinal system in recovery of skilled reaching after cortical stroke: Initial results from a model of ischemic cortical injury. Experimental Brain Research, 233(11), 3231–3251. 10.1007/s00221-015-4390-x

Higo, N. (2021). Non-human Primate Models to Explore the Adaptive Mechanisms After Stroke. Frontiers in Systems Neuroscience, 15. 10.3389/fnsys.2021.760311

Houwink, A., Nijland, R. H., Geurts, A. C., & Kwakkel, G. (2013). Functional Recovery of the Paretic Upper Limb After Stroke: Who Regains Hand Capacity? Archives of Physical Medicine and Rehabilitation, 94(5), 839–844. 10.1016/j.apmr.2012.11.031

Kang, D.-W., Chalela, J. A., Ezzeddine, M. A., & Warach, S. (2003). Association of Ischemic Lesion Patterns on Early Diffusion-Weighted Imaging With TOAST Stroke Subtypes. Archives of Neurology, 60(12), 1730–1734. 10.1001/archneur.60.12.1730

Katan, M., & Luft, A. (2018). Global Burden of Stroke. Seminars in Neurology, 38(2), 208–211. 10.1055/s-0038-1649503

Kosugi, A., Saga, Y., Kudo, M., Koizumi, M., Umeda, T., & Seki, K. (2023). Time course of recovery of different motor functions following a reproducible cortical infarction in non-human primates. Frontiers in Neurology, 14. 10.3389/fneur.2023.1094774

Kwakkel, G., Veerbeek, J. M., Wegen, E. E. H. van, & Wolf, S. L. (2015). Constraint-induced movement therapy after stroke. The Lancet Neurology, 14(2), 224–234. 10.1016/S1474-4422(14)70160-7

Lang, C. E., Bland, M. D., Bailey, R. R., Schaefer, S. Y., & Birkenmeier, R. L. (2013). Assessment of upper extremity impairment, function, and activity after stroke: Foundations for clinical decision making. Journal of Hand Therapy, 26(2), 104–115. 10.1016/j.jht.2012.06.005

Lawrence, D. G., & Kuypers, H. G. (1968). The functional organization of the motor system in the monkey. I. The effects of bilateral pyramidal lesions. Brain: A Journal of Neurology, 91(1), 1–14. 10.1093/brain/91.1.1

Lemon, R. (2004). Cortico-motoneuronal system and dexterous finger movements. Journal of Neurophysiology, 92(6), 3601–3603. 10.1152/jn.00624.2004

Marinković, S., Gibo, H., Milisavljević, M., & Ćetković, M. (2001). Anatomic and clinical correlations of the lenticulostriate arteries. Clinical Anatomy, 14(3), 190–195. 10.1002/ca.1032

Moore, T. L., Bowley, B. G. E., Pessina, M. A., Calderazzo, S. M., Medalla, M., Go, V., Zhang, Z. G., Chopp, M., Finklestein, S., Harbaugh, A. G., Rosene, D. L., & Buller, B. (2019). Mesenchymal derived exosomes enhance recovery of motor function in a monkey model of cortical injury. Restorative Neurology and Neuroscience, 37(4), 347–362. 10.3233/RNN-190910

Moran, D. W., & Schwartz, A. B. (1999). Motor Cortical Representation of Speed and Direction During Reaching. Journal of Neurophysiology, 82(5), 2676–2692. 10.1152/jn.1999.82.5.2676

Morecraft, R. J., Herrick, J. L., Stilwell-Morecraft, K. S., Louie, J. L., Schroeder, C. M., Ottenbacher, J. G., & Schoolfield, M. W. (2002). Localization of arm representation in the corona radiata and internal capsule in the non-human primate. Brain, 125(1), 176–198. 10.1093/brain/awf011

Mugler III, J. P., & Brookeman, J. R. (1991). Rapid three-dimensional T1-weighted MR imaging with the MP-RAGE sequence. Journal of Magnetic Resonance Imaging, 1(5), 561–567. 10.1002/jmri.1880010509

Mugler, J. P., Bao, S., Mulkern, R. V., Guttmann, C. R. G., Robertson, R. L., Jolesz, F. A., & Brookeman, J. R. (2000). Optimized Single-Slab Three-dimensional Spin-Echo MR Imaging of the Brain. Radiology, 216(3), 891–899. 10.1148/radiology.216.3.r00au46891

Murata, Y., & Higo, N. (2016). Development and Characterization of a Macaque Model of Focal Internal Capsular Infarcts. PLOS ONE, 11(5), e0154752. 10.1371/journal.pone.0154752

Nakayama, H., Stig Jørgensen, H., Otto Raaschou, H., & Skyhøj Olsen, T. (1994). Recovery of upper extremity function in stroke patients: The Copenhagen stroke study. Archives of Physical Medicine and Rehabilitation, 75(4), 394–398. 10.1016/0003-9993(94)90161-9

Nudo, R. J., Larson, D., Plautz, E. J., Friel, K. M., Barbay, S., & Frost, S. B. (2003). A Squirrel Monkey Model of Poststroke Motor Recovery. ILAR Journal, 44(2), 161–174. 10.1093/ilar.44.2.161

Nudo, R. J., & Masterton, R. B. (1990). Descending pathways to the spinal cord, III: Sites of origin of the corticospinal tract. Journal of Comparative Neurology, 296(4), 559–583. 10.1002/cne.902960405

Nudo, R. J., Milliken, G. W., Jenkins, W. M., & Merzenich, M. M. (1996). Use-dependent alterations of movement representations in primary motor cortex of adult squirrel monkeys. The Journal of Neuroscience: The Official Journal of the Society for Neuroscience, 16(2), 785–807. 10.1523/JNEUROSCI.16-02-00785.1996

Olivares-Moreno, R., Rodriguez-Moreno, P., Lopez-Virgen, V., Macías, M., Altamira-Camacho, M., & Rojas-Piloni, G. (2021). Corticospinal vs Rubrospinal Revisited: An Evolutionary Perspective for Sensorimotor Integration. Frontiers in Neuroscience, 15. 10.3389/fnins.2021.686481

Orr, C. M. (2017). Locomotor Hand Postures, Carpal Kinematics During Wrist Extension, and Associated Morphology in Anthropoid Primates. The Anatomical Record, 300(2), 382–401. 10.1002/ar.23507

O’Sullivan, S. B., Schmitz, T. J., & Fulk, G. (2019). Physical Rehabilitation. F.A. Davis.

Pearce, T. M., & Moran, D. W. (2012). Strategy-Dependent Encoding of Planned Arm Movements in the Dorsal Premotor Cortex. Science, 337(6097), 984–988. 10.1126/science.1220642

Pizzimenti, M. A., Darling, W. G., Rotella, D. L., McNeal, D. W., Herrick, J. L., Ge, J., Stilwell-Morecraft, K. S., & Morecraft, R. J. (2007). Measurement of Reaching Kinematics and Prehensile Dexterity in Nonhuman Primates. Journal of Neurophysiology, 98(2), 1015–1029. 10.1152/jn.00354.2007

Plautz, E. J., Milliken, G. W., & Nudo, R. J. (2000). Effects of Repetitive Motor Training on Movement Representations in Adult Squirrel Monkeys: Role of Use versus Learning. Neurobiology of Learning and Memory, 74(1), 27–55. 10.1006/nlme.1999.3934

Raffin, E., & Hummel, F. C. (2018). Restoring Motor Functions After Stroke: Multiple Approaches and Opportunities. The Neuroscientist: A Review Journal Bringing Neurobiology, Neurology and Psychiatry, 24(4), 400–416. 10.1177/1073858417737486

Riddle, C. N., Edgley, S. A., & Baker, S. N. (2009). Direct and Indirect Connections with Upper Limb Motoneurons from the Primate Reticulospinal Tract. Journal of Neuroscience, 29(15), 4993–4999. 10.1523/JNEUROSCI.3720-08.2009

Roby-Brami, A., Jarrassé, N., & Parry, R. (2021). Impairment and Compensation in Dexterous Upper-Limb Function After Stroke. From the Direct Consequences of Pyramidal Tract Lesions to Behavioral Involvement of Both Upper-Limbs in Daily Activities. Frontiers in Human Neuroscience, 15. 10.3389/fnhum.2021.662006

Rohlfing, T., Kroenke, C. D., Sullivan, E. V., Dubach, M. F., Bowden, D. M., Grant, K., & Pfefferbaum, A. (2012). The INIA19 Template and NeuroMaps Atlas for Primate Brain Image Parcellation and Spatial Normalization. Frontiers in Neuroinformatics, 6. 10.3389/fninf.2012.00027

Saleem, K. S., & Logothetis, N. K. (2012). A Combined MRI and Histology Atlas of the Rhesus Monkey Brain in Stereotaxic Coordinates. Academic Press.

Sawner, K. A., & LaVigne, J. M. (1992). Brunnstrom’s Movement Therapy in Hemiplegia: A Neurophysiological Approach. Lippincott.

Schalk, G., McFarland, D. J., Hinterberger, T., Birbaumer, N., & Wolpaw, J. R. (2004). BCI2000: A general-purpose brain-computer interface (BCI) system. IEEE Transactions on Biomedical Engineering, 51(6), 1034–1043. 10.1109/TBME.2004.827072

Schiemanck, S. K., Kwakkel, G., Post, M. W. M., Kappelle, L. J., & Prevo, A. J. H. (2008). Impact of internal capsule lesions on outcome of motor hand function at one year post-stroke. Journal of Rehabilitation Medicine, 40(2), 96–101. 10.2340/16501977-0130

Stinear, C. M., Barber, P. A., Smale, P. R., Coxon, J. P., Fleming, M. K., & Byblow, W. D. (2007). Functional potential in chronic stroke patients depends on corticospinal tract integrity. Brain, 130(1), 170–180. 10.1093/brain/awl333

Strick, P. L., Dum, R. P., & Rathelot, J.-A. (2021). The Cortical Motor Areas and the Emergence of Motor Skills: A Neuroanatomical Perspective. Annual Review of Neuroscience, 44(Volume 44, 2021), 425–447. 10.1146/annurev-neuro-070918-050216

Takeuchi, N., & Izumi, S.-I. (2012). Maladaptive Plasticity for Motor Recovery after Stroke: Mechanisms and Approaches. Neural Plasticity, 2012, 359728. 10.1155/2012/359728

Thom, T., Haase, N., Rosamond, W., Howard, V. J., Rumsfeld, J., Manolio, T., Zheng, Z.-J., Flegal, K., O’Donnell, C., Kittner, S., Lloyd-Jones, D., Goff, D. C., Hong, Y., Members of the Statistics Committee and Stroke Statistics Subcommittee, Adams, R., Friday, G., Furie, K., Gorelick, P., Kissela, B., … Writing Group Members. (2006). Heart Disease and Stroke Statistics—2006 Update. Circulation, 113(6), e85–e151. 10.1161/CIRCULATIONAHA.105.171600

Waddell, K. J., Strube, M. J., Bailey, R. R., Klaesner, J. W., Birkenmeier, R. L., Dromerick, A. W., & Lang, C. E. (2017). Does Task-Specific Training Improve Upper Limb Performance in Daily Life Poststroke? Neurorehabilitation and Neural Repair, 31(3), 290–300. 10.1177/1545968316680493

Wang, Y., Liu, G., Hong, D., Chen, F., Ji, X., & Cao, G. (2016). White Matter Injury in Ischemic Stroke. Progress in Neurobiology, 141, 45–60. 10.1016/j.pneurobio.2016.04.005

Wessels, T., Wessels, C., Ellsiepen, A., Reuter, I., Trittmacher, S., Stolz, E., & Jauss, M. (2006). Contribution of Diffusion-Weighted Imaging in Determination of Stroke Etiology. American Journal of Neuroradiology, 27(1), 35–39.

Weston, P., Behr, S., Garosi, L., Maeso, C., & Carrera, I. (2022). Ischemic stroke can have a T1w hyperintense appearance in absence of intralesional hemorrhage. Frontiers in Veterinary Science, 9. 10.3389/fvets.2022.932185

Won, J., Yi, K. S., Choi, C.-H., Jeon, C.-Y., Seo, J., Kim, K., Yeo, H.-G., Park, J., Kim, Y. G., Jin, Y. B., Koo, B.-S., Lim, K. S., Lee, S., Kim, K. J., Choi, W. S., Park, S.-H., Kim, Y.-H., Huh, J.-W., Lee, S.-R., … Lee, Y. (2020). Assessment of Hand Motor Function in a Non-human Primate Model of Ischemic Stroke. Experimental Neurobiology, 29(4), 300–313. 10.5607/en20023

Zhao, B., Shang, G., Chen, J., Geng, X., Ye, X., Xu, G., Wang, J., Zheng, J., Li, H., Akbary, F., Li, S., Lu, J., Ling, F., & Ji, X. (2014). A more consistent intraluminal rhesus monkey model of ischemic stroke. Neural Regeneration Research, 9(23), 2087–2094. 10.4103/1673-5374.147936

Zhi, Y. X., Lukasik, M., Li, M. H., Dolatabadi, E., Wang, R. H., & Taati, B. (2018). Automatic Detection of Compensation During Robotic Stroke Rehabilitation Therapy. IEEE Journal of Translational Engineering in Health and Medicine, 6, 1–7. 10.1109/JTEHM.2017.2780836

